# β4-integrins safeguard nuclear mechanics to suppress prostate cancer progression

**DOI:** 10.64898/2026.09.01.748477

**Authors:** Anette Schmidt, Rong Wei, Paulina Nastaly, Maliha Mehnaz-Enamul, Atharva Agashe, Michal Rychlowski, Katarzyna Bury, Saara Koivusalo, Qin Zhang, Crescenzo Frascogna, Carlo F Natale, Xiayun Yang, Aleksander Jarmakowicz, Anne Ahtikoski, Markku Vaarala, Xiaonan Liu, Katarzyna Ruckemann-Dziurdzinska, Markku Varjosalo, Ewa Bryl, Amrinder Nain, Gong-Hong Wei, Tomasz Wenta, Aki Manninen

## Abstract

Prostate cancer (PCa) progression is accompanied by profound alterations in cell-extracellular matrix (ECM) adhesion, nuclear architecture and mechanical adaptability, yet the molecular mechanisms linking these processes remain poorly understood. Hemidesmosomes (HDs), formed by α6β4-integrins, anchor epithelial cells to the basement membrane and couple extracellular forces to the intermediate filament (IF) cytoskeleton. Here, we identify a previously unrecognized tumor-suppressive function of β4-integrins in preserving nuclear integrity in prostate epithelial cells.

Loss of β4-integrins disrupted the cytokeratin-5 network and its coupling to the nucleus, leading to nuclear softening, lamin remodeling, reduced heterochromatin content and enhanced confined migration. Unexpectedly, proximity-labeling proteomics revealed that β4-integrins engage nuclear pore complex (NPC) components in an α6-independent manner, particularly upon HD disassembly. Selected interactions were validated using proximity ligation and co-immunoprecipitation assays. β4-integrin loss was associated with enlarged nuclear pores and aberrant nucleocytoplasmic transport, including nuclear accumulation of YAP1.

Consistent with these findings, reduced β4-integrin expression in a large PCa tissue cohort correlated with altered nuclear morphology, adverse clinicopathological features, metastatic progression, and poor patient survival. Collectively, our study establishes β4-integrins as a critical molecular link between cell-ECM adhesion, nuclear mechanics and genome integrity.

**SIGNIFICANCE:** β4-integrins interact with nuclear pore complexes revealing nucleoprotective function. Their loss disrupts cytokeratin architecture, alters lamin balance, nuclear transport, and chromatin regulation, providing a mechanism by which β4-integrin loss promotes prostate tumorigenesis.

## INTRODUCTION

Prostate cancer (PCa) is one of the most common malignancies in men with nearly 1.5 million new cases diagnosed annually worldwide(1). Although early detection remains central to effective clinical management, risk stratification and treatment strategies are complicated by the pronounced interpatient heterogeneity at both genetic and phenotypic levels. Current diagnostic strategies rely largely on prostate-specific antigen (PSA) measurements and histopathological evaluation of biopsy specimens. A substantial hereditary component of PCa susceptibility has been firmly established through epidemiological and functional studies, highlighting the need to integrate genetic and molecular determinants into diagnostic strategies(2–8). Clinically heterogeneous intermediate-risk Grade Group 2 (Gleason 3+4) PCa represents a particular diagnostic challenge, as patients in this group carry an increased risk of overtreatment(9). Moreover, the early defining stages of prostate cancer development remain incompletely understood, complicating the distinction between progressive malignant disease and benign proliferation.

Cell-extracellular matrix (ECM) and cell-cell adhesions are central regulators of tissue architecture and mechanical homeostasis, and their dysregulation is a hallmark of cancer progression(10,11). Adhesive structures including focal adhesions (FAs), hemidesmosomes (HDs), adherens junctions (AJs) and desmosomes are mechanically linked to the actin and intermediate filament (IF) cytoskeleton thereby integrating biochemical cues with mechanical forces(12,13). The significance of this mechanochemical coupling is evident from their involvement in a range of pathological conditions, including cancer(14). HDs are α6β4-integrin mediated adhesion complexes that anchor epithelial cells to the basement membrane (BM), a specialized ECM layer underlying epithelia(15). HDs connect via plectin to the keratin IF network, thereby regulating internal mechanical stress and the viscoelastic properties of epithelial cells(16,17). Loss of HDs, a hallmark of the basal-to-luminal transition, is among the most frequent events during cancer initiation and progression, including PCa(18–21). In line with this, we recently reported that HD disassembly in PCa is mainly due to marked reduction of β4-integrin expression across multiple PCa cell lines, independent patient cohorts, and a large tissue microarray (TMA) collection(22).

Importantly, loss of HDs actively contributes to malignant progression as it synergizes with other genomic alterations such as PTEN loss to drive tumorigenesis(22,23). Although depletion of either α6-or β4-integrin subunit disrupts HDs and enhances tumorigenic properties *in vitro*, analyses of patient tissues consistently reveal loss of β4-integrin expression during PCa progression(22). The underlying reason for this selective downregulation remains unexplained. In the absence of β4-integrins, α6-integrins can pair with β1-integrin to form an alternative laminin receptor, α6β1, whereas β4-integrins have not been reported to associate with integrin subunits other than α6. Notably, β4-integrins are expressed in stoichiometric excess relative to α6-subunit and are capable of engaging a broad spectrum of intracellular binding partners independently of α6-integrins(24,25). Moreover, depletion of α6-and β4-integrins elicits distinct cellular phenotypes in PCa models, indicating that each subunit may exert functions beyond the shared role in HD assembly(22).

Together, these observations raise the possibility that β4-integrins exert a specific tumor-suppressive function that is selectively targeted during PCa evolution. To address this hypothesis, we performed a systematic comparative analysis of α6-and β4-integrin knockout phenotypes in normal and malignant prostate epithelial cells. Here, we uncover a previously unrecognized role for β4-integrins in regulating nuclear morphology and mechanics through interactions with nuclear pore components. Loss of β4-integrins was associated with altered nuclear lamina composition, increased nuclear deformability, and enlarged nuclear pores. Importantly, reduced β4-integrin expression and nuclear aberrations correlated with adverse clinical outcomes in PCa patients. Together, our findings establish β4-integrins as critical mediators of epithelial nuclear integrity and mechanotransduction, providing mechanistic insight into why their loss is selectively favored during PCa progression.

## MATERIALS AND METHODS

### Cell culture

RWPE1 and DU145 cell lines were purchased from ATCC and PC3 cell line from Cytion Biosciences. RWPE1 cells were cultured in Keratinocyte SFM medium (Gibco) supplemented with bovine pituitary extract and human recombinant EGF, and standard antibiotics: penicillin (100 units/ml) and streptomycin (100 μg/ml), according to the manufacturer’s protocol. PC3 and DU145 cells were maintained in F12K (Gibco) and MEM (Gibco) medium, respectively containing 10% fetal bovine serum (Gibco) and standard antibiotics. When indicated, 10 µM Y-27632 was added into culture medium. The cells were maintained at 37 °C and 5% CO2. All cell lines were confirmed to be free of mycoplasma during analysis and routinely tested once a month.

### Plasmid construction

The generation of α6-(gRNA: TTTTC TTTGG ACTCA GGGAA AGG targeting exon 6 of *ITGA6*) and β4-integrin (gRNA: CAACT CCATG TCCGA TGATC targeting exon 5 of *ITGB4*)-depleted RWPE1, DU145 and PC3 cells was described in(22). For knock-ins, the HDR-based CRISPR-Cas9 approach was used as described previously(23). To generate GFP-birA-donor plasmid (pUC19-derived) containing the left homology arm of *ITGB4* and right homology arm of *ITGB4* fused with *birA-GFP* sequences, the In-Fusion HD Cloning Kit (Takara Bio, 638910) was applied. The gRNA sequence for *ITGB4* knock-in (ACATGGACCAACAGTTCTTC) was incorporated into plentiCRISPRv2_neo vector as described previously(22). pTRIP-CMV-GFP-FLAG-cGAS E225A-D227A was a gift from Nicolas Manel (Addgene plasmid # 86674; http://n2t.net/addgene:86674; RRID:Addgene_86674)(26). For overexpression of β4-integrin, the sequence of *ITGB4* was cloned into pBabe-puro plasmid. BirA-cDNA was from pcDNA3.1-mycBioID, a gift from Kyle Roux (Addgene plasmid # 35700)(27). For recombinant β4-integrin purification, *ITGB4 G752-G1370* sequence was inserted into pET24b plasmid.

### Viral transduction

Retro-and lentiviral particles were generated as reported in(22). For lentiviral, 5 μg of specific plentiCRISPRv2-gRNA, 3.75 μg psPax2 and 1.25 μg of pVSV-G plasmids were co-transfected into human embryonic kidney 293T cells (ATCC, LGC Standards GmbH, Wesel, Germany; CRL-11268) using Lipofectamine 3000 (Invitrogen, Thermo Fisher Scientific). For retroviral vector generation, 5 μg of pBabe plasmid and 0.5 μg of pVSV-G plasmids were co-transfected into Phoenix-Gag-Pol 293T cells. 1 ml of virus-containing supernatant was filtered using a 0.45 µM spin filter (Sartorius Minisart®), mixed with 4 µg/ml hexadimethrine bromide (Sigma Alrich; H9268) and incubated with 3·10^5^ target cells. Positive clones were selected by the addition of puromycin for at least 7 days or FACS.

### Knock-In Cell Line Generation

For specific β4-integrin knock-in, 1·10^6^ cells were electroporated with 4 µg of *birA-GFP*-donor plasmid in Ingenio® electroporation solution (MIR 50111, Mirus Bio) using Amaxa Nucleofector™ 2b Device (Lonza) and immediately seeded into 5 ml of complete medium in one well of a 6-well plate and incubated at 37°C, 5% CO2 for 6h, as described previously(23). After incubation, non-adherent cells were removed by washing with 1 ml of complete medium followed by transduction or remaining adhered cells with *ITGB4*-targeting gRNA expressing lentiviruses (in lentiCRISPRv2; a gift from Feng Zhang - Addgene plasmid # 52961) as described above(28). After 48h, cells were gently rinsed with complete medium, and antibiotic G418 (Thermo Fisher Scientific) was added for at least 7 days. A minimum of 300 GFP-positive cells were harvested using sorting FACS to generate populations of each variant. Successful knock-in was confirmed by western blotting, fluorescence microscopy and sequencing.

### RNA-Seq

The cells were prepared as detailed in(22). RNA quality was analyzed by using agarose gel electrophoresis, Nanodrop, and RNA Integrity Number (RIN; using Agilent2100). Library construction and sequencing was performed by Novogene Europe (United Kingdom).

### Tissue microarray (TMA)

TMA collection has been previously described in(22). Briefly, it contains a cohort of representative formalin-fixed paraffin-embedded (FFPE) tissue blocks of 221 men operated (radical prostatectomy) from 2005 to 2017 in the Hospital of Oulu under written informed consent with approval of the Regional Ethics Committee of the Northern Ostrobothnia Hospital District (EETTMK 4/2015) in compliance with the Declaration of Helsinki. Two cores from tumor tissue and two cores from benign tissue from each patient were obtained using 2 mm punches. The TMAs were constructed with Galileo TMA CK4500 microarrayer (Isenet, Milan, Italy) in collaboration with the biobank Borealis (Oulu University hospital). The use of pathological archive material was licensed by the National Supervisory Authority for Welfare and Health. This study was approved by the Ethics Council of the Northern Ostrobothnia Hospital District.

### Immunofluorescence staining of cells

The cells grown on 35 mm glass-bottom μ-Dish (IBIDI) were washed twice with PBS and fixed with 4% PFA in PBS for 15 minutes, quenched with 100 mM glycine in PBS for 20 min and then permeabilized with 0.1% Triton X100 in PBS for 15 min. 0.2% gelatin and 0.5% BSA in PBS solution for 1 h has been used before overnight incubation with primary antibody at 4 °C (Supplemental Table S2). After washing 4 times with 0.2% gelatin and 0.5% BSA in PBS, the samples were probed with secondary antibody conjugated with a fluorophore for 1 h or overnight at 4 °C. After the same washing procedure (0.2% gelatin and 0.5% BSA in PBS), the cells were analyzed using Zeiss LSM 780 confocal microscope, Leica SP5, Leica SP8 falcon, Leica SP8X or Zeiss Cell Observer.Z1 Spinning Disc confocal microscope. Colocalization analysis was performed using the ZEN 3.5 (blue edition) and plotting the Pearson’s Correlation Coefficient (PCC, R) from the Colocalization Tool analysis (value is expressed from −1 to +1). Nuclear morphology analysis was done with ImageJ (FIJI) software. Final analysis was performed using GraphPad Prism 9 or 10.

### Immunofluorescence staining of tissues

Validation of TMA staining for β4-integrin was described with details in(22). Briefly, TMA sections were deparaffinized by incubation for 1 h at 55 °C, and antigen retrieval was performed using EnVision FLEX Target Retrieval Solution Low pH (Dako Agilent) in a PT Link device (PT200, Dako Agilent) for 20 min at 97 °C. The TMAs were then incubated overnight at 4 °C with primary antibodies (Supplemental Table S2) prior to incubation with a secondary antibody for 60 min. Nuclei were stained with DAPI, and the sections were mounted with Vectashield Antifade Mounting Medium (Vector Laboratories).

### Quantitative RT-PCR

RNeasy Mini Kit (QIAGEN) was used to extract RNA from cells. RNA purity was analyzed by the 260/280 nm ratio using NanoDrop ONE (Thermo Scientific). RevertAid reverse transcriptase (Thermo Scientific) was used to synthesize cDNA and Power SYBR Green PCR Master Mix (Applied Biosystems) was used for the quantitative RT-PCR analysis. At least three replicates in at least two independent experiments were applied for each gene. All expression data were normalized against GAPDH (control). Primer sequences used in this study are collected in Supplemental Table S3.

### Nanofiber scaffolds

The suspended fibers were spun on hollow scaffolds using the non-electrospinning Spinneret-based tunable engineered parameters (STEP) technique(29–32). For 200 nm and 800 nm fibers, polystyrene with MW: 2,500,000 g/mol (Scientific Polymer Products, CAS no. 1025) was dissolved in xylene (Carolina Chemicals, CAS no. 1330-20-7) for a week. For 2000 nm fibers, polystyrene with MW: 15,000,000 g/mol (Agilent, CAS no. PL2014-9001) was dissolved in xylene (Carolina Chemicals, CAS no. 1330-20-7) for a week. Aligned fibers were extruded through a micropipette (Sutter Instruments) over hollow plastic substrates to form parallel nanofibers with an interfiber spacing of 20 µm to get cells attached to single fibers. Fibers were fused to the substrate with tetrahydrofuran within a custom-made fusing chamber. Nanofiber scaffolds were mounted on 35 mm glass-bottom μ-Dishes (IBIDI) using autoclaved silicon grease and then sterilized with 70% ethanol for 6 min, followed by two rinses with PBS. To ensure cell attachment, scaffolds were coated with 4 μg/ml Human Fibronectin (1918-FN-02M, R&D systems) for 2h at +37°C and finally rinsed with medium before adding the cells. 4·10^5^ cells for RWPE1 and RWPE1 β4-KO, and 2·10^5^ cells PC3 and PC3 β4-KO were seeded and let to attach for 3h (or 24h) in an incubator (37°C, 5% CO2) before fixing and continuing with immunofluorescence staining. Single cells were studied on fibers, excluding those attached to support fibers or multiple fibers. Imaging was performed using a Leica SP8 Falcon confocal microscope with an objective lens of HC PL APO 40x/1.10 W motCORR CS2. Analysis was performed with LAS X (1.4.7.28982) and ImageJ (FIJI) software. 3D rendering and analysis of nuclear staining masks were performed with Imaris 9.2.1 software.

### Invagination analysis

Invaginations caused by fibers were manually measured with Las X (1.4.7.28982) as well as ImageJ (FIJI). In the ImageJ software, fiber and lamin A/C channels were combined to reslice the Z-stack for the YZ direction. A slice showing the deepest invagination caused by the fiber was selected and used for thresholding the lamin A/C channel and creating a solid mask from the nucleus. Lamin A/C shape outline of the selected invagination was used for gathering the invagination curve coordinates. To quantify nuclear invaginations, the manually outlined invaginations were fitted to a bell curve (S = S0e^−y2∕2^*^σ^*^2^) using a custom MATLAB (https://www.mathworks.com/) code to evaluate the individual parameters S0 (invagination depth) and *σ* (invagination spread) as described in(32,33).

### Electron microscopy

Cells were fixed in 2.5% glutaraldehyde (Agar) in PBS buffer for 30 minutes and post-fixed for 1h in 1% osmium tetroxide (Agar). After that time, samples were gradually dehydrated in ethanol, embedded in EPON (Agar), and then cut (65 nm) on ultramicrotome Leica UC7. Uranyless and Reynold’s lead citrate (Delta Microscopies) was used for samples staining. Analyses were performed using Tecnai Spirit BioTWIN transmission electron microscope at 120kV.

### Atomic Force Microscopy

RWPE1 control and RWPE1 β4-KO cells were analyzed using a BioScope Resolve Atomic Force Microscopy (Bruker, Santa Barbara, CA, USA) operating in PeakForce Quantitative Nanomechanical Mapping mode (PF-QNM). 20 × 20 µm glass in 12-well plate were coated with 1 mg/ml collagen I (Merck, CLS354249) for at least 1h at incubator, and then 3·10^5^ cells were seeded on the glass one day before the experiment. Cells were maintained in liquid environment during AFM measurements to preserve their physiological properties. All experiments were conducted at room temperature in a complete medium to maintain cell viability and native mechanical behavior. The PFQNM-LC-V2 probe (Bruker) was used for imaging (nominal resonant frequency f0= 85 kHz; spring constant k = 0.1 N/m, tip radius 70 nm). Images were registered at 512 × 512 pixels with a PeakForce Tapping frequency of 1 kHz and amplitude of 150 nm. Before imaging, the cantilever spring constant was calibrated using the thermal tune method, and the deflection sensitivity was determined on a clean glass surface. Topographic and nanomechanical data were processed and analyzed using NanoScope Analysis software (Bruker). All the data were preprocessed using “Panel fit” and “Flatten” tools with 1^st^ order using NanoScope Analysis 1.8 and the extracted data were analyzed using Gwyddion 2.67 software.

### Design and fabrication of cell confinement device

The cell confinement device was designed using a 3D CAD software. The model was exported as an STL file and imported into Describe (Nanoscribe GmbH) to define the printing parameters. A parametric sweep was performed to optimize fabrication conditions in terms of resolution and geometric fidelity. The master was fabricated using a Nanoscribe Photonic Professional GT system equipped with a 25× objective. IP-S photoresist was used as the printing material. To improve the adhesion between the polymerized IP-S master and the substrate and to prevent delamination during PDMS demolding, a silanization protocol was carried out prior to printing. The substrates were first cleaned and then activated using an oxygen plasma treatment to render the surface hydrophilic. A silane solution was freshly prepared by mixing 50 mL ethanol with 250 µL of 3-(trimethoxysilyl)propyl methacrylate (CAS 2530-85-0), followed by the addition of 1.5 mL of diluted acetic acid (1:10 glacial acetic acid:water) immediately before use. The substrates were submerged in the silane solution for at least 10 minutes. After treatment, the substrates were rinsed with isopropanol and gently dried with a nitrogen stream. IP-S photoresist was deposited directly onto the silanized ITO substrate, which was mounted on the holder for dip-in laser lithography. Following laser writing, the printed sample was immersed in mr-Dev (Micro Resist Technology) for 20 minutes to remove unpolymerized resin. The substrate was then transferred into a second beaker containing isopropanol to wash away residual developer. The developed master was used as a negative mold for device fabrication via soft lithography. PDMS (Sylgard 184, Dow Corning; 10:1 base-to-curing agent ratio) was poured onto the master and degassed under vacuum to eliminate trapped air bubbles. The sample was cured in an oven at 80 °C for 2 hours. After curing, the PDMS layer was carefully peeled off from the master and individual devices were cut from the slab. For microscopic imaging and cell experiments, the PDMS devices were irreversibly bonded to glass coverslips. A second oxygen plasma treatment (50 W, 1 minute) was applied to both the PDMS and the glass substrate to activate their surfaces. Immediately after plasma exposure, the two surfaces were brought into conformal contact to achieve permanent bonding suitable for imaging and fluidic sealing. The micro-channel layout was designed using CAD software. The device features three inlet ports for cell loading, which lead into a rectangular entry region with a height of 50 µm. Within this area, supporting micro-pillars were integrated to prevent roof collapse during PDMS-to-glass bonding, while still allowing cells to move freely between them and reach the micro-channel interface. Two types of microchannels were designed: i) non-constrictive channels, with dimensions of 8 µm × 8 µm (height × width) and a length of 200 µm for analysis of basic migration of the cells and ii) constrictive channels, featuring a narrowing (striction) down to 4 µm in the XY plane, while maintaining a constant height of 8 µm in Z. These geometries were chosen to enable controlled confinement and passage of cells under different mechanical constraints.

### Western blotting

Cells at 80–90% confluence were washed in DPBS (Gibco) and scraped in RIPA buffer: 10 mM Tris-HCl pH 8.0, 150 mM NaCl, 0.5% SDS, 1% IGEPAL, 1% sodium deoxycholate containing 2 mM PMSF (phenylmethylsulfonyl fluoride), 10 μg/ml aprotinin, and 10 μg/ml leupeptin. Protein concentration was assessed by BCA Protein Assay Kit (Pierce). Thirty µg of protein lysate was resolved by SDS-PAGE and transferred onto a Protran pure 0.2 micron nitrocellulose (Perkin Elmer). After 1h of incubation in 5% skimmed milk, the membranes were probed with primary antibodies (Supplemental Table S2) overnight at 4 °C. The bands were visualized using Lumi-Light Western Blotting Substrate (Roche) and Azure Biosystems Inc., Bioanalytical Imaging System, model c400.

### Nuclei isolation

Adherent cells were washed once with DPBS, trypsinized and collected by centrifugation at 1100 rpm for 3 min. The cell pellet was resuspended in 1 mL DPBS, centrifuged again under the same conditions and then resuspended in 0.5 ml hypotonic buffer containing 0.33 M sucrose, 10 mM HEPES pH 7.4, 1 mM MgCl₂, 10 mM NaCl and 0.1% IGEPAL CA-630. The suspension was centrifuged at 3000 rpm for 10 min at 4 °C. This washing step was repeated three times. The supernatants from the first two centrifugation steps were collected as the cytoplasmic fraction. The remaining pellet was incubated on ice for 30 minutes in the same buffer containing additionally 0.1% Tween-20 and 1 % BSA, and then washed twice more under the same conditions. Finally, the isolated nuclei were resuspended in 1% BSA in PBS with RNA inhibitor for further microscopic analysis or immediately lysed in RIPA buffer for western blotting analysis.

### Biotin Ligation and Sample Preparation for LC-MS/MS

Cells were seeded as 4.5 × 10^4^/cm on three 15 cm Ø dishes and cultured for 24 h in the presence of 50 μm biotin as described in(25). Cells were washed three times with cold PBS+/+, scraped and immediately frozen with liquid nitrogen. Cysteine bonds were reduced with 5 mm Tris (2-carboxyethyl)phosphine (TCEP) (Merck) for 20 min at 37°C and alkylated with 10 mm iodoacetamide (Merck) for 20 min at room temperature in the dark. After overnight digestion with a total of 1 μg of Sequencing Grade Modified Trypsin (Promega, Madison, Wisconsin), the samples were quenched with 10% trifluoroacetic acid (TFA) and purified with C-18 Micro SpinColumns (Nest Group Inc., Thermo Fischer Scientific, Helsinki, Finland) in 0.1% TFA in 50% acetonitrile (ACN). Final samples were reconstituted in 30 μl 0.1% TFA and 1% ACN in LC-MS grade water.

### Liquid chromatography-mass spectrometry (LC-MS/MS)

LC-MS/MS analyses were performed using an Evosep One LC system coupled to a timsTOF Pro mass spectrometer (Bruker) via a CaptiveSpray source. Peptide separation was achieved on an Evosep column (8 cm × 150 µm, 1.5 µm C18 beads) using a standardized “60 samples per day (60SPD)” gradient (Buffer A: 0.1% formic acid in water; Buffer B: 0.1% formic acid in acetonitrile).

MS data were acquired in positive ion DDA mode with parallel accumulation serial fragmentation (PASEF), performing 10 PASEF scans per topN cycle. The resulting raw files were processed using MSFragger against the human UniProtKB database, with all instrument and label-free quantification parameters left at their default settings. Cysteine carbamidomethylation was set as a fixed modification, while methionine oxidation and lysine/N-terminal biotinylation were set as variable modifications.

### Recombinant protein purification

The *E. coli* BL21(DE3) clpP::cmR cells transformed with pET24_*ITGB4* plasmid were used to overproduce β4-integrin (aa 752-1370) with the His6 tag, in pET System (Novagen, San Diego, CA, USA). The recombinant proteins were purified by affinity chromatography using the Co-NTA resin (Clontech Inc.) followed by gel filtration chromatography (Superdex 200, Sigma-Aldrich). The purity of the recombinant proteins used for further studies was estimated to be more than 95% as judged by SDS-polyacrylamide gel electrophoresis. The recombinant human NUP98 (aa 1-880) was purchased from BIOMATIK, USA (RPC27839).

### Pull down

β4-integrin (aa G752-G1370) recombinant protein was dialyzed against the pull-down buffer 50 mM Tris-HCl pH 7.5, 150 mM NaCl, 1% Triton X-100, 10% glycerol and 1 x protease inhibitor (Roche 04693159001). The PC3 cells were homogenized in ice-cold pull-down buffer and the lysates were cleared by centrifugation at 14000g for 30 min at 4 °C followed by incubation with Co-NTA beads. After that, 2 mg of lysate proteins were incubated with the tagged β4-integrin protein (0.04 mg) overnight at 4 °C. The Co-NTA beads were then added to the mixture and incubated for next 3 h at 4 °C. The beads were recovered by centrifugation and washed five times with 500 μl of the pull-down buffer. Protein complexes were eluted by using 50 µl of 2x SDS sample loading buffer and incubation at 95°C for 10 minutes.

### Co-immunoprecipitation (Co-IP)

Cells growing on two 10 cm Ø plates (∼80 % confluency) were washed twice with DPBS and harvested by scraping in 0.5 ml IP cold buffer (50 mM Tris-HCl pH 7.5, 150 mM NaCl, 1% Triton X-100, 10% glycerol, 1 mM EDTA and 1 x protease inhibitor (Roche 04693159001)). After a 30-minute incubation on ice, during which the cells were vortexed every 5 minutes, the samples were sonicated (Q800R sonicator, Q Sonica) at 4°C water bath for 30 sec on 100% amplitude and centrifuged for 30 minutes at 14000g at 4°C. Supernatant was pre-cleared by incubation with 50 µl Protein G magnetic beads (Invitrogen 10004) for 1h at 4°C on a gentle shaker and then incubated overnight at 4°C with 5-8 µg of antibody. The next day, 50 µl of G-protein magnetic beads was added and the mixture was gently agitated for a additional 6 hours at 4°C. The complexes were collected using a magnetic rack, following five 5-minute washes with 500 µl of IP buffer, using 30 µl of 2x SDS sample loading buffer and incubation at 95°C for 10 minutes.

### Proximity ligation assay

For visualization of the β4-integrin complexes with α6-integrin, NUP98 and TPR in cells, the proximity ligation assay (PLA) (Duolink® PLA Technology based on Duolink® In Situ Red Starter Kit Mouse/Rabbit – Merck DUO92101) was applied, as described in (34). RWPE1 and RWPE1 α6-KO cells were seeded at the density of 2 · 10^5^ cells per well in 12-well plate on collagen I-coated glass cover slips and allowed to grow overnight. The cells were fixed in 4% formaldehyde, for 20 min at room temperature, incubated with 200 mM glycine for 20 min and then permeabilized by adding 0.1% Triton-X100 in PBS for 7 min at 4°C. To limit the non-specific interactions the cells were incubated in Blocking solution for 1 h at 37°C and followed to the manufacturer’s protocol with the reagents and media provided in the PLA kit (Merck). The reaction with the primary antibodies: mouse anti-β4-integrin (1:50) and rabbit: anti-α6-integrin (1:200), anti-NUP98 (1:100) or rabbit anti-β4-integrin (1:50) and mouse anti-TPR (1:100) was carried out for 2 h at room temperature. After washing, the incubation with the secondary oligonucleotide-conjugated antibodies: anti-rabbit PLA plus and anti-mouse PLA minus probes was conducted for 1 h at 37 °C in a humidity chamber. All following steps (ligation, amplification and mounting) were performed according to the manufacturer’s protocol. Immunofluorescence signals were analyzed under a Leica microscope (TCS SP8).

### Enzyme-linked immunosorbent assay (ELISA)

The assay was performed as described previously in(34). Briefly, the 96-well plates were coated with 0.4 μg recombinant β4-integrin or BSA in PBS at 4°C overnight and then washed three times in EL buffer (2% BSA, 0.05% Tween-20 in PBS), blocked for 2h in the same buffer containing 0.1% Tween20 and incubated with increased amount (0–8 μg) of NUP98 protein (1-880aa, RPC27839, BIOMATIK, USA) resuspended in EL buffer for 2h at 37°C. The anti-NUP98 antibody diluted in EL buffer (1:1000) were used as the primary antibodies for 2h at RT. The anti-mouse-conjugated IgG (1:7000) were used as the secondary antibodies. The reaction was stopped by the addition of 1 M H2SO4. The optical density was measured at 450 nm using Perkin Elmer EnSpire multimode plate reader. At least two independent experiments with at least three technical repeats were performed.

### Data filtering steps and analysis

Rstudio (version: 4.4.3) was used for Gene Ontology (GO) and Gene set enrichment analysis (GSEA) of both RNA-Seq and interactome data. Normalization and log2 conversion were carried out to identify the differentially expressed genes (DEGs), and the DEGs are displayed as volcano plots. The cut off conditions were as |log2-fold change| ≥ 1 and adjusted *P-*value (adj. *P*) < 0.05 in RNAseq data, while for differentially expressed proteins (DEPs) were set as |log2-fold change| ≥ 1 and *P-*value < 0.05. The Venn diagram tool (https://bioinfogp.cnb.csic.es/tools/venny/index.html) was used to compare and analyze the results of the intersection analysis. GO and GSEA Enrichment analyses were conducted to determine whether a series of a priori-defined biological processes were enriched with adj *P* < 0.05 chosen for further analysis. Clustering analysis of common DEPs were identified in α6-KO and PTEN/α6-dKO groups using ‘GseaVis’ package (Version: 0.1.0)(35). Potential candidates were identified using sankeygoplot in ‘GseaVis’ package.

Protein–protein interaction confidence scores were calculated using the Mass spectrometry Interaction STatistics (MiST) algorithm as described in(36). For each bait–prey pair, MiST integrates three quantitative parameters: reproducibility (R) across biological replicates, specificity (S) of prey enrichment across all baits, and abundance (A) based on normalized spectral counts or intensity values. The final MiST score was computed according to 0.40*A + 0.40*R + 0.20*S representing a slightly modified weighting scheme compared with the original publication, where higher MiST scores indicate more confident interactions. Interactions with MiST ≥ 0.55, R ≥ 0.6, and S ≥ 0.4 were considered high-confidence candidates. To further categorize interactors, we defined core complex components (MiST ≥ 0.55, R ≥ 0.6, freq ≥ 3) to keep the highly-expressed proteins in each group such as ITGB4. MiST calculations and visualization were performed in R (v4.4.3) using a customized pipeline based on the public MiST implementation (https://github.com/everschueren/mist).

High-confidence protein-protein interactions from the protein interactome mass spectrometry data were identified statistically using the Significance Analysis of INTeractome (SAINT)-express tool (version 3.6.0)(37) and the Contaminant Repository for Affinity Purification (CRAPome)(38). Eight BioID runs targeting GFP were used as negative controls. Proteins were considered as HCIs if they met all the following criteria: a SAINT express probability score ≥ 0.78 (Bayesian FDR < 0.05) and a CRAPome frequency < 20%.

All protein-protein interaction networks were generated using string database (https://string-db.org/)(39) and visualized in Cytoscape software(40).

Kaplan-Meier curves (95% CI) were computed using Prism software (ver. 9.5.1), for the ITGB4 analysis the median values were used as a cut-off. The statistical significance and p-value were calculated using Log-rank (Mantel-Cox) test. Cox proportional hazards model was applied to calculate the hazard ratio (HR) for analysis the relative risk between different patient groups.

### Statistical analysis

*Data are presented as box-and-whisker plots (min to max values)* of at least three independent experiments, unless otherwise indicated in the figure legend. The single points in the box plots represent the number of analyzed objects or repeats. Comparative data were analyzed with the unpaired t-test, One-way or Two-way ANOVA or nonparametric assay using GraphPad Prism 9/10 software. The results with p-value lower than 0.05 (*), 0.01 (**), 0.001 (***) or 0.0001 (****) were considered statistically significant.

## RESULTS

### Transcriptomic profiling reveals both shared and subunit-specific consequences of α6-and β4-integrin loss

Formation of the hemidesmosomal (HD) adhesions is a well-established function of α6β4-integrins(41). Previous studies have linked the loss of functional α6β4-integrins to altered cell adhesion dynamics and increased migratory activity, with selective downregulation of the β4-subunit being a recurrent feature in PCa patient samples(23,42). To elucidate molecular differences that might underlie the preferential loss of β4-integrins during tumor progression, we performed transcriptomic RNA sequencing of parental RWPE1 prostate epithelial cells (control), RWPE1 α6-integrin-knockout (α6-KO) cells and RWPE1 β4-KO cells.

Compared with the control cells, α6-KO cells showed a total of 1723 differentially expressed genes (DEGs), including 655 upregulated and 1068 downregulated transcripts (Fig 1A). In comparison, β4-KO cells displayed a broader transcriptional perturbation, with 2155 DEGs, including 937 upregulated and 1218 downregulated genes (Fig 1B). More than 40% (1118 genes) of these DEGs were shared between α6-KO and β4-KO cells (Fig 1C). As expected, shared DEGs were significantly enriched for pathways related to cell adhesion and ECM-related categories (Supplemental Table S1). However, unbiased Gene Ontology (GO) analysis revealed that the most prominently enriched shared categories were associated with nuclear and cell-cycle–regulation (Fig 1D).

**Figure 1.**
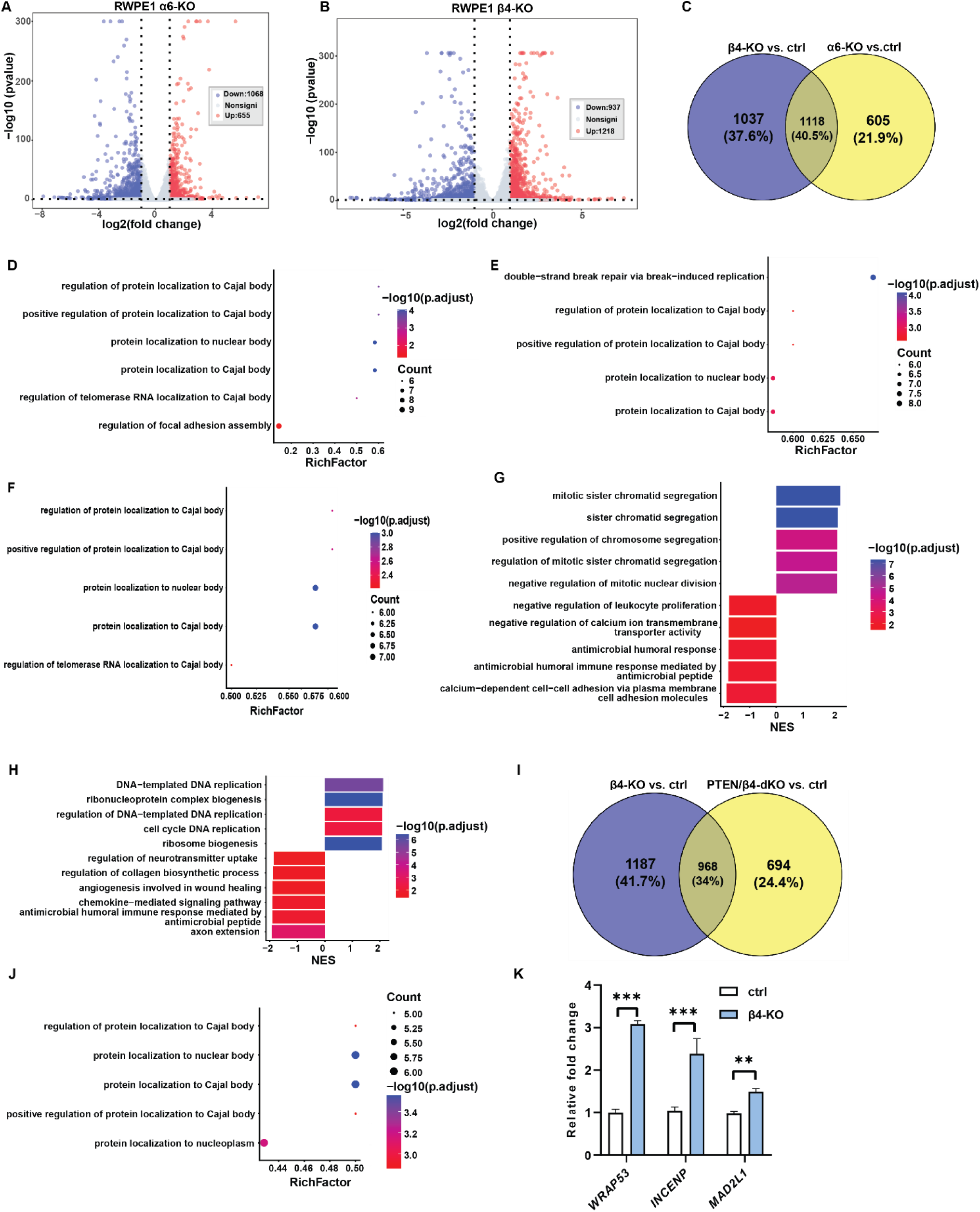
Loss of β4-integrins affects nucleus-related transcriptional programs. **A-B)** Volcano plots of DEGs identified in RWPE1 α6-KO vs. control (ctrl) cells (A) and β4-KO vs. control cells (B). A total of 1723 DEGs out of 16068 detected genes were identified in α6-KO cells compared with control, including 655 up-and 1068 downregulated genes. In β4-KO cells, 2155 DEGs out of 16709 detected genes were identified, including 937 up-and 1218 downregulated genes. Red and blue dots indicate significantly up-and downregulated genes, respectively. Significance was defined as |log₂ (fold change)| > 1 and adjusted p < 0.05. **C)** Venn diagram showing overlap of DEGs between RWPE1 β4-KO vs. control and RWPE1 α6-KO vs. control cells. A total of 1118 common DEGs were identified, with 1037 DEGs specific to β4-KO and 605 DEGs specific to α6-KO. **D-F)** GO enrichment analysis of the shared DEGs (D), β4-KO-specific DEGs (E), and α6-KO-specific DEGs (F). The x-axis represents the Rich factor (ratio of DEGs number to the total number of genes annotated to a given GO term), and the y-axis lists the top 5 enriched biological process terms. Dot size corresponds to the number of genes involved in each term, and bar color indicates significance level (-log₁₀ adjusted p-value). **G-H)** GSEA summary plots of genes ranked by log₂ (fold change) in RWPE1 β4-KO vs. control (G) and RWPE1 α6-KO vs. control (H). The x-axis represents the normalized enrichment score (NES), and the y-axis lists the top 5 significantly up-and downregulated biological process terms. Bar color corresponds to the significance level (-log₁₀ adjusted p-value). **I)** Venn diagram showing common DEGs between RWPE1 β4-KO vs. control and RWPE1 PTEN/β4-dKO vs. control cells. A total of 968 shared DEGs were identified. **J)** GO enrichment analysis of the 968 shared DEGs identified in β4-KO and PTEN/β4-dKO cells. Axes and plot parameters as described in panels D-F. **K)** qPCR validation of selected DEGs from the most enriched pathways. Data are presented as mean ± SD from at least three technical replicates from at least two biological repeats. Full list of enriched pathways shown in panels D-H and J is provided in Supplemental Table S1.

A substantial set of subunit-biased transcriptional changes was also detected: 1037 DEGs were more prominent in β4-KO cells, and 605 DEGs were preferentially associated with α6-KO cells (Fig 1C). GO analysis revealed that nuclear processes, particularly mitosis and Cajal body–associated functions, were most strongly enriched in β4-KO cells, although these pathways were also significantly represented in α6-KO cells (Fig 1E-F). Gene Set Enrichment Analysis (GSEA) further highlighted subunit-specific differences. α6-KO cells displayed stronger upregulation of biosynthetic programs, including ribosome biogenesis and rRNA metabolism, DNA replication, and metabolic activity. In contrast, β4-KO cells showed more pronounced upregulation of chromatin-and mitosis-related pathways, accompanied by downregulation of gene silencing and cell–cell adhesion pathways (Fig 1G-H).

In RWPE1 cells, loss of either α6-or β4-integrin subunit leads to strong downregulation of the remaining partner(22). Therefore, the extensive overlap between α6-KO and β4-KO transcriptomes is at least partly explained by this reciprocal regulation, complicating efforts to assign the observed effects specifically to either subunit. Recently, we showed that concomitant PTEN loss uncouples this interdependency, as PTEN/β4-double knockout (dKO) cells retain robust α6-integrin expression(22). We therefore compared transcriptomes of RWPE1 β4-KO and RWPE1 PTEN/β4-dKO cells (Fig 1I). This analysis identified 968 shared DEGs enriched pathways consistent with those identified in the β4-KO dataset (Fig 1J). Quantitative PCR validation of selected DEGs from the most significantly enriched pathways confirmed these transcriptional changes (Fig 1K). Parallel analysis of PTEN/α6-dKO cells demonstrated preservation of the biosynthetic and ribosomal programs characteristic of α6-KO cells (Supplemental Fig S1A-B). By contrast, PTEN-KO cells displayed distinct immune and inflammation-dominated transcriptional profiles (Supplemental Fig S1C-D).

Collectively, these data demonstrate that depletion of either α6-or β4-integrin perturbs cell adhesion and ECM-related transcriptional programs and induces cell-cycle associated responses. Importantly, loss of β4-integrins elicited a disproportionately strong nuclear and chromatin-centered transcriptional responses, supporting the notion that β4-integrins perform functions beyond HD formation, potentially linked to regulation of nuclear organization.

### β4-KO cells display altered nuclear morphology

To study the impact of β4-integrin depletion on nuclear architecture, we performed confocal microscopy with lamin A/C and DAPI staining, complemented by electron microscopy analysis. In RWPE1, DU145, and PC3 cells, loss of β4-integrins consistently resulted in nuclei with increased cross-sectional area and pronounced alterations in the nuclear envelope (NE) morphology compared with control cells (Fig 2; Supplemental Figs. S2A–G and S3A–G). Quantitative shape analysis of maximum intensity projections from confocal imaging and nuclear cross-sections obtained by confocal and electron microscopy, showed that nuclei of β4-deficient cells appeared larger and exhibited increased folding and invagination of the NE relative to nuclei in control cells. These structural changes were reflected in altered nuclear shape metrics, including reduced circularity, solidity, and roundness, as well as an increased aspect ratio (Fig 2; Supplemental Figs. S2A-G and S3A-G). Similar observations were found in breast cancer cell line with β4-integrin depletion (Supplemental Fig S4A-F) and RWPE1 with β4-integrin overexpression (Supplemental Figs. S4G-L). Collectively, these findings establish that β4-integrin loss alone is sufficient to drive consistent and marked alterations in nuclear morphology across non-transformed and malignant prostate epithelial cells, in line with the transcriptomic evidence linking β4-integrins to the control of nuclear organization.

**Figure 2.**
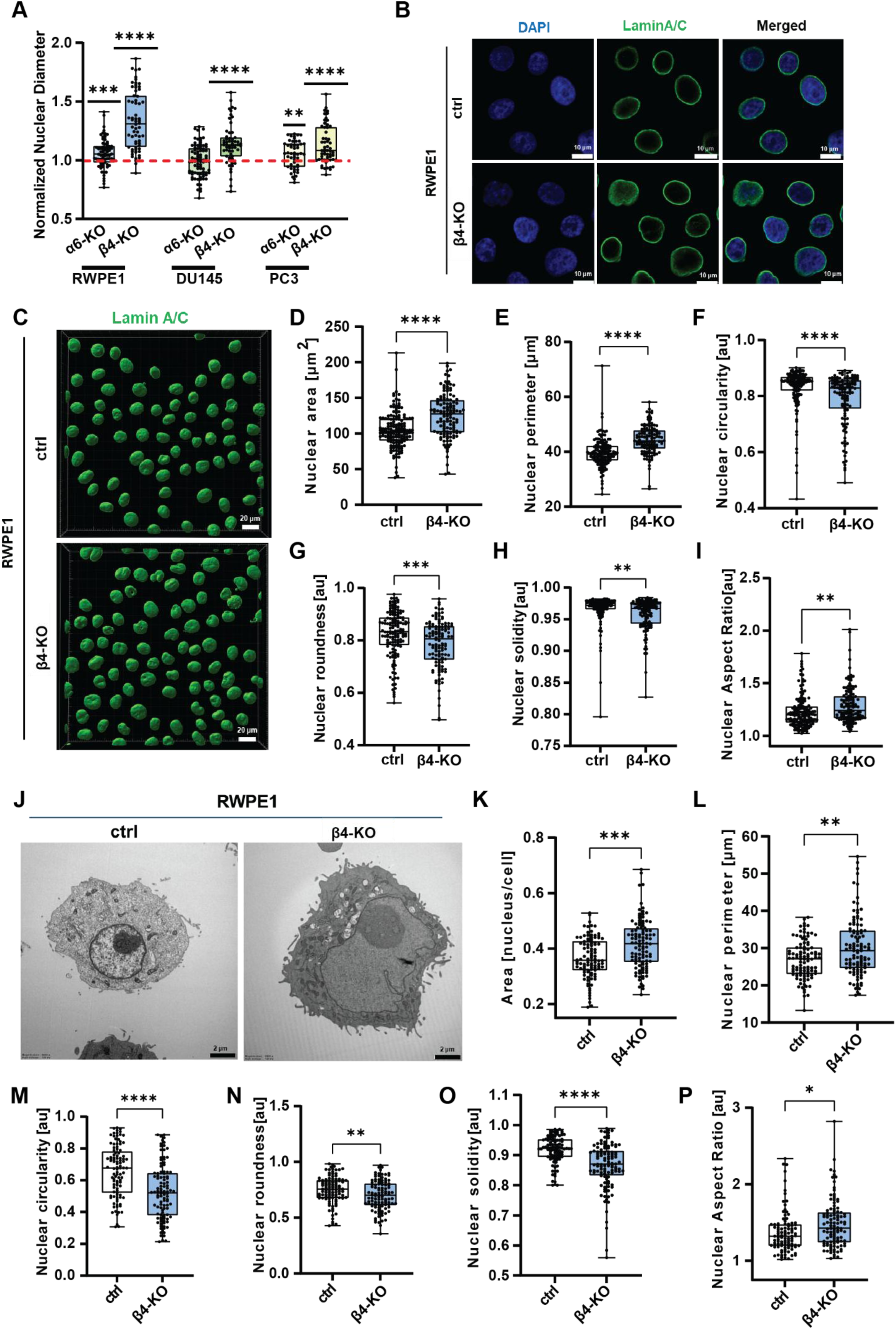
Loss of β4-integrins alters nuclear size and shape. **A)** Quantification of nuclear diameter in RWPE1, DU145, and PC3 parental cells and their α6-KO or β4-KO variants. Values were normalized to the mean nuclear diameter of the corresponding parental control cells. The red dashed line indicates the control mean. **B)** Confocal images of RWPE1 control and β4-KO cells stained for DAPI (blue) and lamin A/C (green). Images were acquired using a Leica SP8 Falcon microscope with a 40× water objective and analyzed using ImageJ (FIJI). Scale bars, 10 µm. **C)** 3D-rendered lamin A/C staining visualizing nuclear morphology, generated using Imaris software. Scale bars, 20 µm. **D-I)** Quantitative nuclear-shape analysis from maximum-intensity Z-stack projections of lamin A/C–and DAPI-stained cells. Parameters measured using ImageJ (FIJI): nuclear area (D), perimeter (E), circularity (F), roundness (G), solidity (H), and aspect ratio (I). **J)** Representative electron microscopy images of RWPE1 control and RWPE1 β4-KO nuclei. Analysis was performed using Tecnai Spirit BioTWIN transmission electron microscope. Scale bars, 2 µm. **K-P)** Nuclear-shape quantification from electron microscopy images, measured using ImageJ (FIJI): nuclear area (K), perimeter (L), circularity (M), roundness (N), solidity (O), and aspect ratio (P). Data are presented as box-and-whisker plots (min to max). Each dot represents an individual nucleus.

### Loss of β4-integrins disrupts cytokeratin-5 anchorage and weakens IF-nucleus mechanical coupling

The nucleus is mechanically coupled to both the endoplasmic reticulum and the cytoskeleton via the Linker of Nucleoskeleton and Cytoskeleton (LINC) complex(43). Integrin adhesion complexes at the cell–ECM interface are critical initiators of mechanotransduction, transmitting extracellular forces into the cell interior. While much emphasis has been placed on studies addressing the role of FAs and the actin cytoskeleton in nuclear mechanotransduction, less is known about the role of HDs linking the ECM to the IF cytoskeleton to provide viscoelastic support(44,45). β4-integrins recruit plectin to tether cytokeratin-5 (CK5), the major IF component of basal epithelial cells, thereby stabilizing the IF network and linking it to the NE(46). This arrangement suggests that IFs serve as important transmitters of HDs-mediated mechanical signals to the nucleus.

To investigate how loss of β4-integrins affects the IF cytoskeleton, we analyzed CK5 organization. Total internal reflection fluorescence (TIRF) microscopy revealed robust basal CK5 staining in confluent RWPE1 and PC3 cells. In contrast, β4-KO cells displayed markedly reduced basal CK5 localization, indicating defective anchoring of CK5 at the basal surface (Fig 3A-B). Furthermore, whereas CK5 filaments in control cells formed a distinct perinuclear cage encircling the nucleus, this organization was disrupted in β4-KO cells, suggesting impaired IF-NE connectivity (Fig 3C-D).

**Figure 3.**
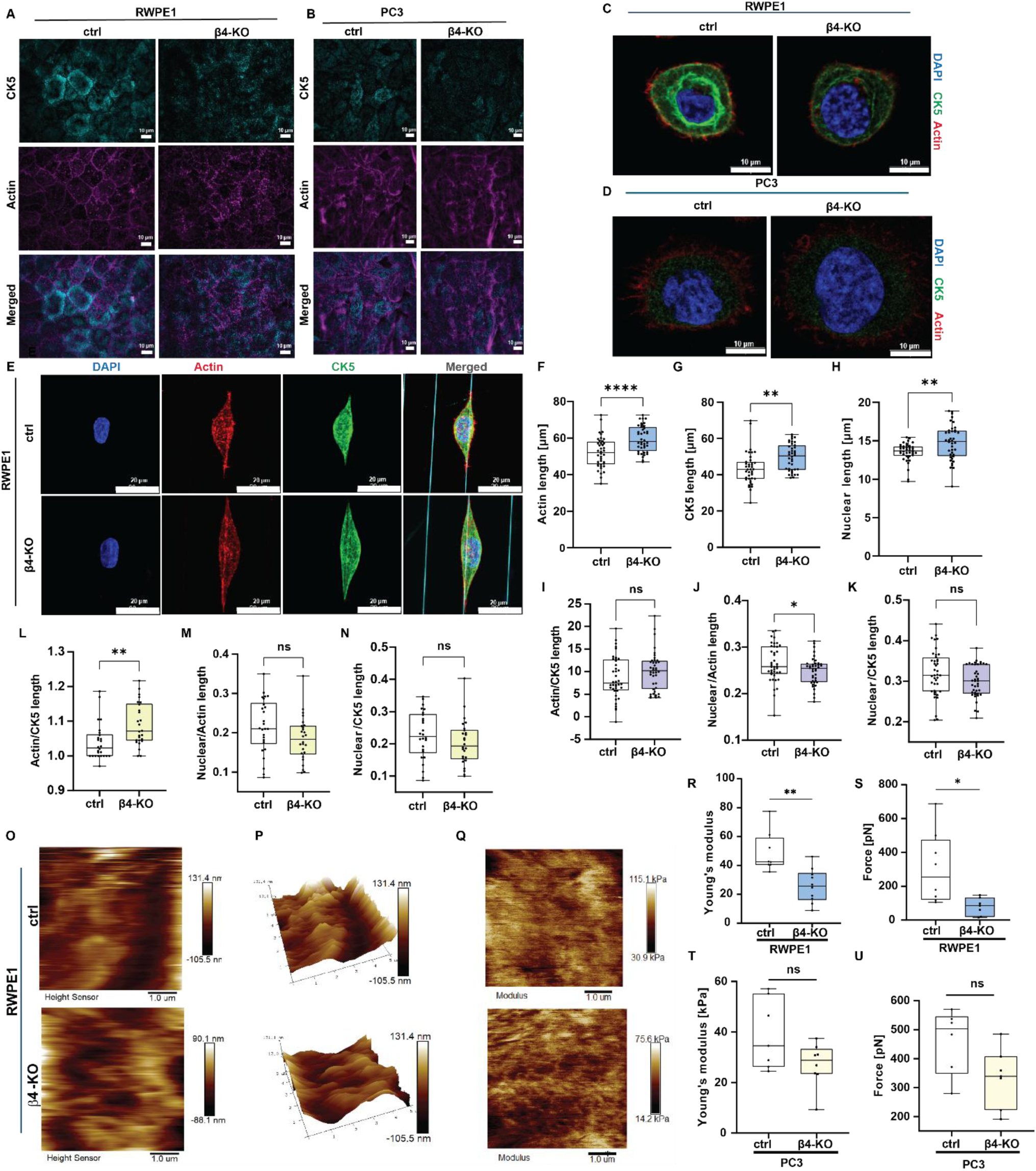
Loss of β4 integrins disrupts CK5 anchorage and weakens the mechanical coupling between the IF network and the nucleus. **A-B)** Total internal reflection fluorescence (TIRF) imaging of cytokeratin 5 (CK5, cyan) and actin (magenta) in RWPE1 (A) and PC3 (B) control and β4-KO cells. Images were acquired with a Zeiss Cell Observer.Z1 spinning-disk confocal microscope using a 63× oil objective. Scale bars, 10 µm. **C-D)** Confocal z-stack images of non-confluent RWPE1 (C) and PC3 (D) cells stained for CK5 (green), actin (red), and DAPI (blue), imaged 3 h after seeding using a Leica SP8 Falcon microscope with a 40× water objective. Scale bars, 10 µm. **E)** Maximum-intensity projections of CK5 (green), actin (red), and DAPI (blue) staining in RWPE1 cells cultured on aligned 800-nm nanofibers. Images were acquired using Leica SP8 Falcon confocal microscope with a 40x water objective. **F-H)** Quantification of actin (F), CK5 (G), and nucleus length (H) in RWPE1 control and β4-KO cells grown on nanofibers **I-K)** Quantification of actin/CK5 length ratio (I), nucleus/actin length ratio (J), and nucleus/CK5 length ratio (K) in RWPE1 control and β4-KO cells grown on nanofibers. **L-N)** Corresponding analysis as shown in panel I-K, performed in PC3 control and β4-KO cells. **O-Q)** Representative AFM height-sensor images and 3D topography maps of RWPE1 control and β4-KO cells. Cells were analyzed using a BioScope Resolve Atomic Force Microscopy (Bruker, Santa Barbara, CA, USA) operating in PeakForce Quantitative Nanomechanical Mapping mode (PF-QNM). **R-U)** AFM-based measurements of cell stiffness and adhesion. Young’s modulus (R, S) and adhesion force (T, U) were quantified in RWPE1 and PC3 cells, respectively. Data are presented as box-and-whisker plots (min to max). Each dot represents an individual measurement.

Cellular mechanotransduction plays a critical role in how cells respond to diverse microenvironmental stresses, such as altered stiffness, crowding and fibrillar matrix organization, all of which are characteristic features of tumor lesions(47–49). At the invasive front of PCa epithelium, cells are exposed to fibrillar stromal ECM that imposes anisotropic mechanical tension. To model these physiological mechanical constraints *in vitro*, we cultured cells on aligned nanofiber arrays that reproduce the topology and directional tension cues of the in vivo ECM(50). Analysis of actin, CK5, and nuclear organization in nanofiber-grown cells showed that the actin cytoskeleton extended further toward the cell periphery than the CK5-positive IF network (Fig 3E-H). By quantifying the relative lengths of the actin and CK5 networks and correlating them with nuclear length in control and β4-KO RWPE1 and PC3 cells, we found that the actin/CK5 ratio increased, and the nucleus/actin ratio decreased in β4-KO cells (Fig 3I–N). Notably, the nucleus/CK5 ratio remained unchanged (Fig 3K, N), suggesting that nuclear elongation in β4-KO cells becomes at least partially uncoupled from overall cell shape.

Because CK5 contributes to epithelial stiffness and protects cells from mechanical stress, loss of HD-mediated anchoring of the CK5 network may reduce cellular stiffness and promote a more aggressive phenotype(51). To test this, we measured cell elasticity using atomic force microscopy (AFM). Depletion of β4 integrins significantly reduced cell stiffness in both RWPE1 and PC3 cells, as indicated by lower Young’s modulus values (Fig 3R, T). It also decreased the tip-cell detachment force measured during AFM retraction (Fig 3S, U), consistent with reduced membrane tension and membrane-cytoskeleton coupling. These mechanical defects are consistent with the disrupted CK5 network in β4-KO cells, which reduces membrane-cytoskeleton coupling and weakens cellular mechanical support.

Together, these findings indicate that β4-integrin loss disrupts basal anchoring of CK5 IF network, leading to destabilization of the IF cytoskeleton and impaired mechanical coupling between the IF cytoskeleton and the nucleus. These defects contribute to the aberrant nuclear morphology and altered nuclear mechanics observed in β4-deficient cells.

### β4-integrin depletion leads to nuclear softening

An altered nuclear morphology is often linked to changes in nuclear mechanics(52). Because nuclear shape is strongly influenced by mechanical inputs transmitted through the cytoskeleton and cell-ECM adhesions, we next examined how β4-integrin depletion affects nuclear behavior in a fibrillar environment. For this purpose, we used the nanofiber system described above, which provides defined mechanical cues through fiber diameter and alignment(32,50).

Consistent with our 2D observations, nuclei in β4-KO cells displayed more ruffled and irregular lamin A/C staining compared with the smooth, elliptical nuclei of control cells (Fig 4A). These abnormalities appeared even more pronounced on nanofibers (Fig 4A, C). Control nuclei aligned along the fiber axis, whereas β4-KO nuclei showed less consistent alignment. On thicker fibers (2000 nm and 800 nm), β4-KO nuclei were significantly longer than control nuclei (Fig 4A-D), likely reflecting enhanced FA dynamics in β4-KO cells, since thicker fibers support FA maturation(23). Fiber-induced nuclear invaginations were strongly diameter-dependent and were more pronounced in β4-KO cells, which also exhibited increased nuclear folding (Fig 4A, C). To test whether these nuclear changes were influenced by actomyosin-generated tension, we treated cells with the ROCK inhibitor Y-27632. ROCK inhibition reduced nuclear elongation on fibers and induced pronounced NE ruffling in both cell types, with stronger effects in β4-KO cells (Fig 4B-D).

**Figure 4.**
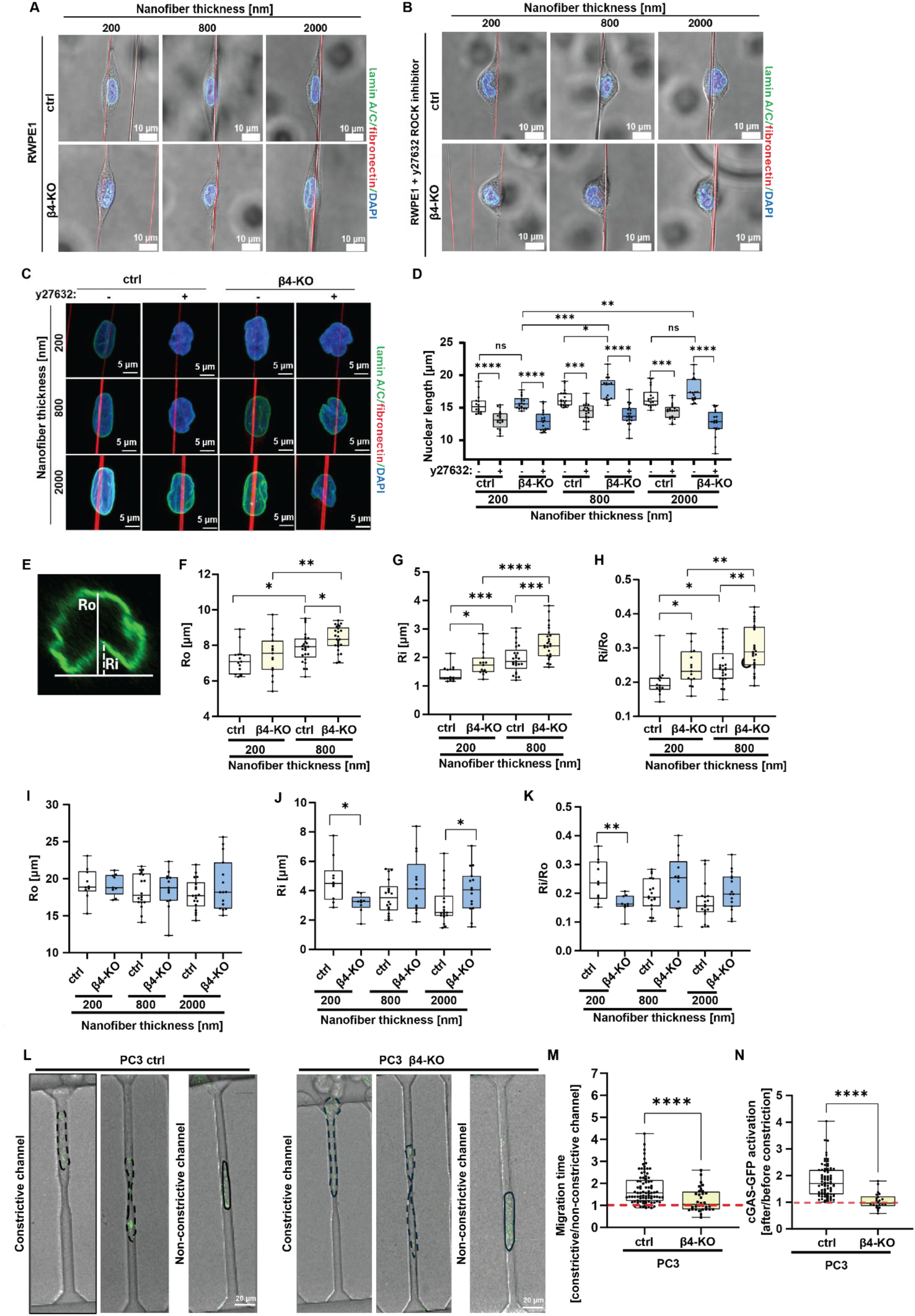
Loss of β4-integrin leads to nuclear softening. **A-B)** Merged images of RWPE1 control and β4-KO cells on fibronectin-coated nanofibers of indicated thickness. Cells were visualized in brightfield and stained with DAPI (blue) and lamin A/C (green), with nanofibers shown in red. Cells were cultured in complete K-SFM without (A) or with the ROCK inhibitor Y-27632 (B). Brightfield images show overall cell shape. Images were acquired using Leica SP8 Falcon confocal microscope with a 40x water objective. Scale bar, 10 µm. **C)** Maximum intensity projections of nuclei in RWPE1 control and β4-KO cells grown on nanofibers of different thicknesses. Lamin A/C (green) outlines the nuclear envelope (NE) and DAPI (blue) marks the chromatin. Images were acquired using Leica SP8 Falcon confocal microscope with a 40x water objective. **D)** Quantification of nuclear length in RWPE1 control and β4-KO cells across nanofiber diameters of 200 nm, 800 nm, and 2000 nm, with and without Y-27632 treatment. **E)** yz-side view of a lamin A/C–stained nucleus illustrating how invagination parameters were measured. Ro indicates the overall nuclear diameter and Ri indicates the depth of nuclear envelope invagination. Lamin A/C staining was used to outline the nuclear envelope. **F-H)** Quantification of nuclear invagination parameters in PC3 control and β4-KO cells grown on nanofibers of 200 nm and 800 nm thickness. Parameters include Ro (F), Ri (G), and the Ri/Ro ratio (H). **I-K)** Quantification of nuclear invagination parameters, Ro (I), Ri (J), and Ri/Ro ratio (K), in RWPE1 control and β4-KO cells grown on nanofibers of 200 nm, 800 nm, and 2000 nm thickness. All analyses in panels F-K were imaged using a Leica SP8 Falcon confocal microscope with a 40x water objective and quantified with LAS X software (1.4.7.28982). **L)** Analysis of nuclear mechanical behavior using a 3D-fabricated cell confinement device. PC3 control and β4-KO cells expressing cGAS-GFP were introduced into channels containing both constrictive and non-constrictive segments. Migration through non-constrictive channels was used to control for differences in basal migration ability. Solid black lines outline cell boundaries, and dashed black lines indicate nuclear contours. Representative time-lapse images correspond to Supplemental Videos 1-4. Images were acquired using Leica SP8X microscopy with a 20x oil objective. **M)** Comparison of the time required for PC3 control and β4-KO cells to pass through constrictive channels relative to non-constrictive channels. **N)** Quantification of cGAS-GFP activation following nuclear constriction. Activation was calculated as the ratio of normalized cGAS-GFP fluorescence intensity after versus before migration through the constrictive segment. Data in all panels are presented as box-and-whisker plots (min to max). Each dot represents a single cell.

To quantify nuclear mechanics, we measured NE curvature by calculating the ratio of invagination depth (Ri) to nuclear height (Ro) (Fig 4E). This local deformation metric reflects the viscoelastic properties of the NE and integrates contributions from cytoskeletal integrity, actomyosin contractility, and the ability of cells to form mature adhesions on fibers of different diameters. In metastatic PC3 cells grown for 3h on 200 and 800 nm fibers, β4-KO nuclei showed increased NE curvature relative to controls, suggesting that β4-deficient PC3 nuclei were softer (Fig 4F-K). In RWPE1 cells, β4-KO nuclei were softer on 800 nm fibers, whereas on 200 nm fibers they showed shallower invaginations, suggesting increased NE tension (Fig 4I-K). On 2000 nm fibers, RWPE1 β4-KO nuclei again appeared softer (Fig 4I-K). These differences likely reflect the distinct adhesive capacities of RWPE1 and PC3 cells on thin fibers, with RWPE1 cells having reduced ability to form mature FAs on 200 nm fibers.

Nuclear softening facilitates cancer cell migration through confined spaces during metastasis(53). To test whether loss of β4-integrins alters confined migration, we used a 3D-fabricated device containing non-constrictive (8µm, control migration) and constrictive (4µm) channels (Fig 4L). PC3 β4-KO cells required significantly less time than control cells to migrate through the constrictive channels, indicating increased nuclear deformability (Fig 4M). Moreover, β4-KO cells showed significantly lower cGAS-GFP foci formation after constriction, consistent with fewer NE rupture events (Fig 4N). Together, our data is consistent with a model in which loss of β4-integrins leads to nuclear softening, increased deformability, and improved capacity to migrate through confined spaces (Fig 4L-N).

### Reduced levels of β4-integrin expression in PCa tissues are associated with altered nuclear properties and adverse clinical outcome

To assess the physiological and clinical relevance of β4-integrin-dependent nuclear alterations, we analyzed nuclear morphology in relation to β4-integrin expression in a tissue microarray (TMA) comprising paired normal and tumor samples from 221 PCa patients. Reduced β4-integrin expression correlated with increased nuclear area and perimeter and decreased nuclear eccentricity, indicating larger nuclei with a less elongated but more irregular contour, morphological features that have been associated with aggressive cancers and adverse clinical outcomes (Fig 5A-G)(54,55).

**Figure 5.**
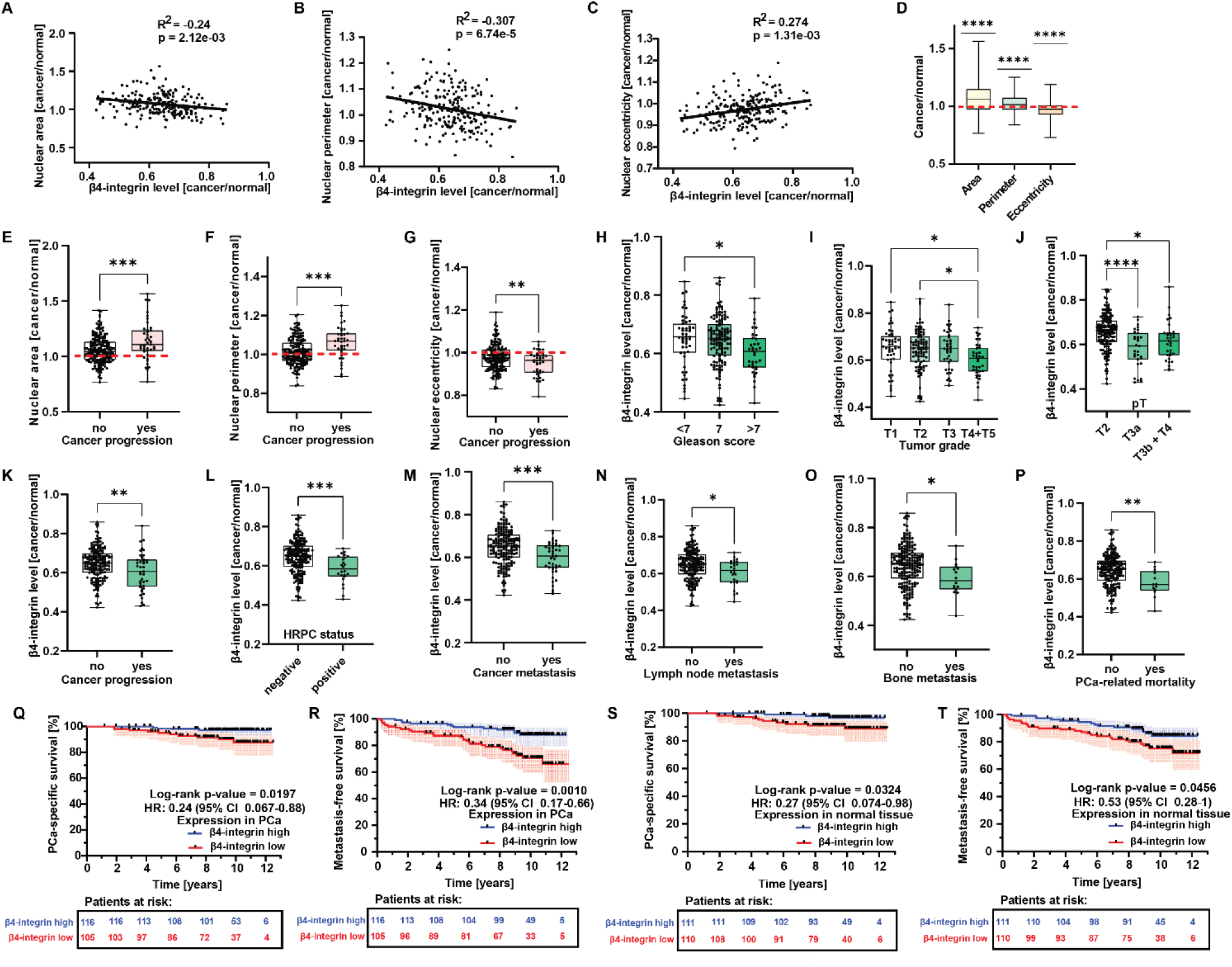
Reduced β4-integrin expression correlates with altered nuclear morphology and adverse clinical outcome in PCa. **A-C)** Correlation between β4-integrin expression (cancer-to-normal ratio per patient) and nuclear morphometric parameters in the TMA cohort of 221 patients. Parameters include nuclear area (A), nuclear perimeter (B), and nuclear eccentricity (C). **D)** Comparison of nuclear morphology features in matched normal and tumor tissue from the same patients, showing increased nuclear area and perimeter and decreased eccentricity in tumor nuclei. **E-G)** Association of nuclear morphology parameters with clinicopathological progression. Increased nuclear area (E) and perimeter (F), and reduced nuclear eccentricity (G), were associated with higher-grade or more advanced PCa. **H-P)** Association between β4-integrin expression and clinicopathological parameters. Lower β4-integrin levels correlated with higher Gleason score (H), tumor grade (I), pathological stage (J), advanced disease status (K), hormone-refractory status (L), and presence of metastases (M), including lymph node (N) and bone metastases (O), as well as PCa-specific mortality (P). All β4-integrin values are presented as cancer-to-normal ratios for each patient. Each point represents one patient. Box-and-whisker plots show minimum to maximum values. **Q-T)** Kaplan-Meier curves showing PCa-specific survival (Q) and metastasis-free survival (R) stratified by β4-integrin expression in tumor tissue, and corresponding analyses for phenotypically normal prostate tissue (S-T). Log-rank test, shaded regions represent 95% confidence intervals. Statistical significance is indicated as follows: ns, p ≥ 0.05; * p < 0.05; ** p < 0.01; *** p < 0.001; **** p < 0.0001.

Importantly, lower β4-integrin levels also correlated with multiple clinicopathological indicators of PCa severity, including higher Gleason score (Fig 5H), increased tumor grade (Fig 5I), advanced pathological stage (Fig 5J), progression to advanced disease (Fig 5K), hormone-refractory status (Fig 5L) and presence of metastases (Fig 5M), including both lymph node (Fig 5N) and bone metastases (Fig 5O). In addition, reduced β4-integrin expression was significantly associated with PCa-related mortality (Fig 5P).

Kaplan-Meier analysis revealed that patients with low β4-integrin expression had significantly shorter PCa-specific survival (Fig 5Q) and metastasis-free survival (Fig 5R). Notably, similar associations were observed when β4-integrin expression was measured in phenotypically normal epithelium from the same patients (Fig 5S-T), suggesting that loss of β4-integrin expression may be an early event in PCa progression.

Taken together, these data indicate that downregulation of β4-integrins is linked to aberrant nuclear morphology and adverse clinical outcome in PCa.

### Loss of β4-integrins induces a lamin isoform switch

Lamins are nuclear IF proteins that provide structural and mechanical support to the nucleus and maintain nuclear shape. A-type lamins (lamin A and lamin C) are enriched in differentiated cells and contribute to nuclear mechanics, chromatin organization, transcriptional regulation and cell cycle control(56). B-type lamins (lamin B1 and lamin B2) are constitutively expressed in almost all cell types and support nuclear lamina (NL) integrity and genome maintenance(57,58). The relative abundance of A-and B-type lamins influences NL properties and contributes to mechanical behavior of the nucleus.

To examine whether β4-integrin loss affects lamin composition, we analyzed lamin B1 and lamin A/C protein levels in RWPE1 and PC3 cells. Western blot analysis showed a consistent increase in lamin B1 and a reduction in lamin A/C levels in β4-KO cells (Fig 6A-D), suggesting that loss of β4-integrins shifts the balance toward a more lamin B1-dominant NL.

**Figure 6.**
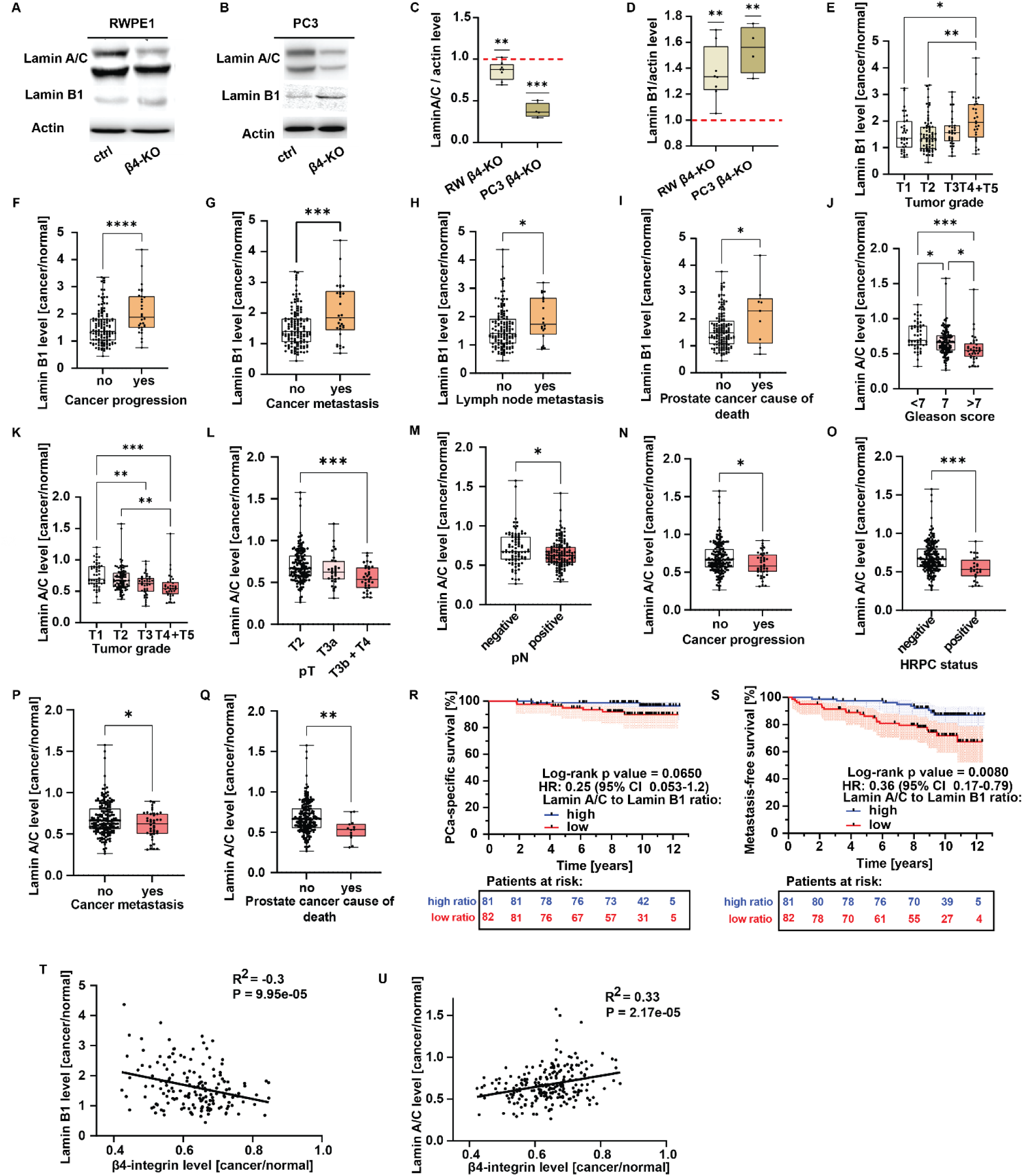
Loss of β4-integrins alters NL composition by shifting lamin isoform levels. **A-B)** Western blot analysis of lamin B1 and lamin A/C expression in RWPE1 (A) and PC3 (B) control and β4-KO cells. **C-D)** Quantification of lamin A/C (C) and lamin B1 (D) protein levels. Red dashed lines indicate the normalized mean values of the corresponding parental control cells. Each dot represents a biological replicate. Data are shown as box-and-whisker plots (min to max). **E-I)** Association of lamin B1 levels with clinicopathological variables in the TMA cohort of 221 men with PCa. Higher lamin B1 expression (cancer-to-normal ratio) correlated with higher tumor grade (E), PCa progression (F), metastasis (G), lymph node metastasis (H), and PCa-related mortality (I). **J-Q)** Association of lamin A/C levels with clinicopathological variables in the same TMA cohort. Lower lamin A/C levels correlated with higher Gleason score (J), tumor grade (K), pathological stage (L), lymph node involvement (M), PCa progression (N), hormone-refractory status (O), metastasis (P) and PCa-related mortality (Q). Values are shown as cancer-to-normal ratios per patient. **R-S)** Kaplan-Meier plots showing PCa-specific survival (R) and metastasis-free survival (S) stratified by the lamin A/C to lamin B1 ratio. Log-rank test, shaded areas indicate 95% confidence intervals. **T-U**) Correlation between β4-integrin expression (cancer-to-normal ratio) and lamin B1 (T) or lamin A/C (U) levels across the 221 PCa patients in the TMA cohort. Each dot represents one patient.

To assess the clinical relevance of this lamin switch, we analyzed lamin expression in paired normal and tumor samples from the same 221-patients PCa TMA cohort. Elevated lamin B1 levels were associated with higher tumor grade, disease progression, metastasis, lymph node involvement and PCa-related mortality (Fig 6E-I). Conversely, reduced lamin A/C expression correlated with higher Gleason score, advanced pathological stage, lymph node metastasis, PCa progression, hormone-refractory disease, metastasis, and cancer-related death (Fig 6J-Q).

Because the lamin A/C to lamin B1 ratio reflects NL composition and correlates with nuclear mechanical properties, we examined its association with clinical outcome. A lower lamin A/C to lamin B1 ratio was associated with significantly shorter PCa-specific survival (Fig 6R) and metastasis-free survival (Fig 6S). Importantly, lamin expression levels correlated with β4-integrin expression across the TMA cohort. Lamin B1 showed a negative correlation, whereas lamin A/C exhibited a positive correlation with β4-integrin levels (Fig 6T-U).

Together, these findings indicate that reduction of β4-integrins, leading to HD disassembly, is linked to altered NL composition characterized by increased lamin B1 and decreased lamin A/C. In PCa tumor tissues these properties correlate with poor clinical outcome.

### β4-integrin interacts with nuclear pore complex proteins

Our data suggest that β4-integrin may have a tumor-suppressive role in PCa linked to their ability to preserve nuclear integrity. To define the underlying molecular mechanisms, we analyzed the β4-integrin interactome in prostate epithelial cells. For this objective, we overexpressed β4-integrin fused to BirA biotin ligase in RWPE1 and PC3 cell lines. Control cells and cells with overexpression of N-myristoylated BirA-GFP treated with biotin were used as controls. In both cell lines exogenously expressed β4-integrin was found to interact with several HD components including α6-integrin, plectin, and collagen XVII (Supplemental Fig S5). Intriguingly, GO analysis of high-confidence interactors (HCIs) identified using MIST algorithm not only showed enrichment of pathways associated with cell adhesion, but also revealed potential β4-integrin interactions with several nuclear proteins (Supplemental Fig S5).

Previous studies have indicated that β4-integrins are typically expressed in stoichiometric excess relative to α6-integrins(24) and can engage with multiple interaction partners in the absence of functional HDs(25). However, whether β4-subunits possess HD-independent cellular functions has remained unclear. Thus, to ensure efficient capture of both HD-associated and HD-independent interaction partners, we characterized the β4-interactome in HD-forming control cells and in HD-deficient α6-KO cells. For this specific purpose, we generated a BioID proximity-labeling system by inserting the BirA biotin ligase sequence in-frame with GFP into the endogenous *ITGB4* locus using homology directed repair (HDR)-mediated CRISPR–Cas9 method (23). The GFP tag facilitated clone selection, and a flexible linker containing a myc epitope and six glycine residues minimized interference with β4-integrin binding interactions. This strategy enabled proximity-dependent biotinylation of proteins interacting with β4-integrin expressed from the endogenous locus (Supplemental Fig S6A-B).

In control cells with intact HDs, BioID identified robust interactions between β4-integrins and canonical HD components (Supplemental Fig S6C-G, J, L, N). Upon HD disassembly in α6-KO cells, β4-integrin proximity interactome became enriched in a distinct set of candidate partners (Supplemental Fig S4H-I, K, M, O). In total, 415 differentially enriched proteins (DEPs) were identified (194 gained and 221 decreased or lost; Supplemental Fig S4C). GO analysis revealed significant enrichment of pathways related to desmosome organization, nuclear pore assembly and actin nucleation (Supplemental Fig S4D). Consistently, GSEA showed that β4-integrin preferentially associated with proteins involved in nuclear pore function when HDs were absent (Supplemental Fig S4E), suggesting a potential contribution to nuclear envelope homeostasis.

Given our previous observation of functional synergy between HD loss and PTEN depletion, we extended this analysis to PTEN/α6-dKO cells. Comparative analysis of α6-KO and PTEN/α6-dKO cells identified 255 shared β4-interacting proteins (Fig 7A). Hierarchical clustering segregated these interactors into two major groups: (i) 170 DEPs that were lost from the β4-integrin interactome upon HD disassembly, and (ii) 85 DEPs gained in α6-KO and PTEN/α6-dKO cells (Fig 7B). GO analysis associated the first cluster primarily with cell–cell and cell–matrix adhesion (Fig 7C), consistent with the established role of β4-integrins in HDs. In contrast, the second cluster was strongly associated with nuclear pore organization and nucleocytoplasmic transport (Fig 7D).

**Figure 7.**
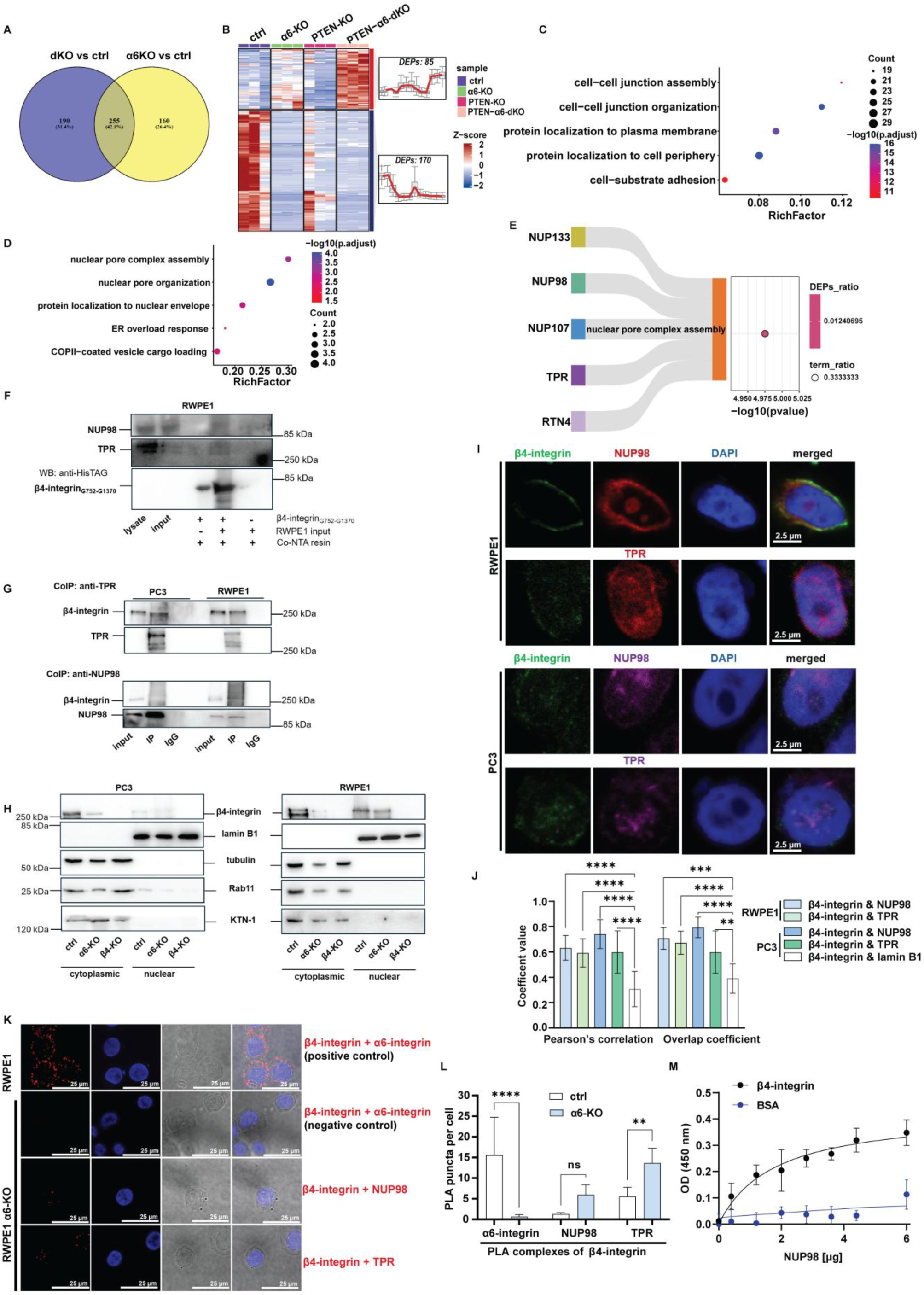
β4-integrins interact with NPC proteins and these interactions are enhanced upon α6-integrin depletion. **A-B)** Venn diagram (A) and hierarchical clustering heatmap (B) of DEPs shared between α6-KO vs control and PTEN/α6-dKO vs control cells. Protein abundance values are shown as Z-scores. Two major clusters containing 85 and 170 DEPs, respectively, are highlighted with representative mean-expression profiles. **C-D)** GO enrichment analysis of the two DEP clusters shown in panel B. Cluster of 170 DEPs (C) represents β4-integrin interacting proteins lost in α6-KO cells, whereas the cluster of 85 DEPs (D) represents β4-interacting proteins gained upon α6-integrin depletion. The x-axis shows the Rich factor and the y-axis lists the top enriched biological processes. Dot size corresponds to the number of proteins in each term and color indicates-log₁₀(adjusted p-value). **E)** Sankey plot showing candidate NPC proteins enriched in the top pathway “nuclear pore complex assembly” and identified as potential β4-integrin interactors upon α6-integrin depletion. Candidates include NUP133, NUP98, NUP107, TPR, and RTN4. **F)** Pull down assay results of recombinant β4-integrin (G752-G1370 aa) incubated with RWPE1 cell lysate. The lysate was pre-cleared by incubation with Co-NTA beads prior incubation with β4-integrin. **G)** Co-immunoprecipitation (Co-IP) results of complexes NUP98 & β4-integrin and TPR & β4-integrin in RWPE1 and DU145 cells. IgG control serves as a negative control. **H)** Western blotting analysis of isolated nuclei from RWPE1 and PC3 showing nuclear localization of β4-integrin. Efficiency of isolation was verified by analysis of lamin B1 as a nuclear marker, tubulin – cytoplasmic marker, KTN-1 - endoplasmic reticulum marker and Rab11 – marker of recycling endosomes. **I)** Immunofluorescence analysis of the β4-integrin localization in the nuclei of normal (RWPE1) and cancer PC3 cells. **J)** Quantification of colocalization of β4-integrin with NUP98 and TPR in isolated nuclei from RWPE1 and PC3 cell lines. The colocalization was assessed by analysis of Pearson correlation and Overlap coefficients using LASX software. Lamin B1 serves as a negative control beceause this protein was not identified as a potential β4-integrin partner. Data are presented as mean ± SD. Asterisks indicate significance (p-value: *<0.05; **<0.01; ***<0.001). **K)** Proximity ligation assay (PLA) showing interaction of β4-integrin with α6-integrin (positive control), NUP98, and TPR in RWPE1 control and α6-KO cells. The interaction of α6-integrin with β4-integrin in α6-KO cells was included as a negative control. Images were acquired using Leica SP8X microscopy with a 63x oil objective. Scale bar, 25 µm. **L)** Quantification of PLA puncta per cell. Analysis was performed using ImageJ (FIJI). **M)** Validation of direct interaction between β4-integrin and NUP98 performed by using ELISA assay. The plate was coated with recombinant β4-integrin and incubated with increasing amount of recombinant NUP98. BSA serves a negative control. The full list of the enriched pathways shown in panels B-C and F-G is provided in Supplemental Table S1.

To validate these findings, we applied two complementary confidence-scoring frameworks, MiST and SAINT, which rely on distinct statistical principles to identify high-confidence proximity interactors. In control cells, β4-integrin preferentially associated with classical HD proteins and components of the cell-cell and cell-matrix adhesion machineries (Supplemental Fig. S4F, J). Disruption of HDs by α6-integrin depletion resulted in a pronounced shift in the β4-integrin proximity landscape. Both MiST-and SAINT-based analyses revealed reduction of adhesion-related partners and enrichment of proteins linked to nuclear pore organization and protein localization to the nucleus (Supplemental Fig. S4H-I, K, M).

This shift was also observed in PTEN/α6-dKO cells, indicating that HD disassembly promotes acquisition of nuclear-associated β4-integrin partners also in cells with metastatic potential. Network and pathway enrichment analyses highlighted nucleoporins including NUP98, and TPR as prominent candidates gained upon HD disruption (MiST scores 0.689 and 0.608, respectively; Fig. 7E and Supplemental Fig. S4H-I, K, M, O). These proteins constitute structural and regulatory modules of the nuclear pore complex(59), raising the possibility that β4-integrins influence NPC organization and function.

Next, we validated these potential interactions using a pull down assay, co-immunoprecipitation (Co-IP), proximity ligation assays (PLA) and immunofluorescence microscopy. For pull down assay we purified recombinant β4-integrin (G752-G1370) protein as the bait and incubated that with RWPE1 and PC3 cell lysates. NUP98 was specifically captured by recombinant β4-integrin-containing beads (Figure 7F). TPR signal was somewhat weaker in β4-integrin-containing samples. This could be due to observed non-specific binding of TPR to the Co-NTA resin, that will result in significant reduction the cellular pool of TPR after the pre-clearing step with Co-NTA resin only (Fig 7F). To further confirm these protein-protein interactions, we performed Co-IP experiments in RWPE1 and PC3 cells, in which endogenous NUP98 and TPR were immunoprecipitated with the anti-NUP98 and anti-TPR antibodies, respectively. β4-integrins co-precipitated with both NUP98 and TPR (Fig 7G). However, the levels of β4-integrin co-precipitating with NUP98 were relatively low in PC3 cells (Fig 7G).

β4-integrins are considered to exert their functions at the plasma membrane. To characterize possible nuclear functions, we studied whether β4-integrins can be observed in the proximity of the NE-associated proteins, NUP98 and TPR. To this end, we isolated nuclei and analyzed them by western blotting. Cytoplasmic and nuclear fraction were verified by analysis of lamin B1 as a nuclear marker, tubulin (cytoplasmic marker), KTN-1 (endoplasmic reticulum marker) and Rab11 (recycling endosome marker). Curiously, clearly detectable signal of β4-integrins was found in the nuclear fraction of RWPE1 and PC3 cells (Fig 7H). Next, we stained isolated nuclei to study the nuclear localization of β4-integrin in more detail. Immunofluorescence analysis in intact cells demonstrated that the majority of β4-integrin signal is at the plasma membrane or its proximity thereby complicating the verification of lower levels of potential nuclear localization. However, analysis in isolated nuclei, revealed β4-integrin localization at the NE, where it colocalized with NUP98 and TPR (Fig 7I-J). Lamin B1, is a nuclear protein located in close proximity of NE but which was not among potential β4-interacting proteins, was used as a negative control that did not show significant co-localization with β4-integrin (Fig 7J). Moreover, we used PLA to independently validate and visualize β4-integrin interactions with NUP98 and TPR. The well-characterized interaction between β4-and α6-integrins served as a positive control in parental cells and as a negative control in α6-knockout cells, validating assay specificity (Fig 7K-L). PLA analysis confirmed that NUP98 and TPR formed detectable complexes with β4-integrins (Fig 7K-L). In HD-deficient α6-KO cells, these interactions were found to be even more prominent (Fig 7K-L).

To examine if β4-integrin directly interacts with NUP98, we performed an ELISA assay using recombinant proteins. A 96-well plate was coated with recombinant β4-integrin, followed by incubation with increasing concentrations of purified recombinant NUP98. As shown in Fig 7M, the OD value in β4-integrin-containing wells increased upon addition of NUP98 in a dose-dependent manner, whilst no effect was seen in wells coated with BSA. Collectively, all these findings strongly support a mechanism where β4-integrins interact with nuclear pore proteins. Moreover, these interactions appear to become even more prominent upon HD disassembly.

Subsequently, to assess the functional relevance of the β4-integrin interactome data, we analyzed nuclear pore morphology by transmission electron microscopy. Strikingly, nuclear pore diameter and perimeter were significantly increased in β4-integrin-depleted RWPE1-, DU145-, and PC3 β4-KO cells compared with controls (Fig 8A-F and Supplemental Fig S3H-J). To examine whether these structural alterations might influence nucleocytoplasmic trafficking, we analyzed the subcellular localization of the mechanosensitive transcriptional regulator YAP1, which dynamically shuttles between the cytoplasm and nucleus through the NPC. RWPE1 β4-KO cells displayed a pronounced nuclear accumulation of YAP compared with control cells (Fig 8G-H), consistent with altered NPC function. Significant albeit less prominent nuclear pore enlargement was also observed in RWPE1 α6-KO cells (Fig 8C-D), which are known to downregulate β4-integrin levels(22). However, in contrast to β4-KO cells, RWPE1 α6-KO cells did not exhibit increased nuclear YAP accumulation (Fig. 8H). This suggests that although reduced β4-integrin levels may influence nuclear pore size, a critical threshold of β4-integrin loss must be exceeded before NPC permeability is sufficiently compromised to permit aberrant nuclear entry of YAP.

**Figure 8.**
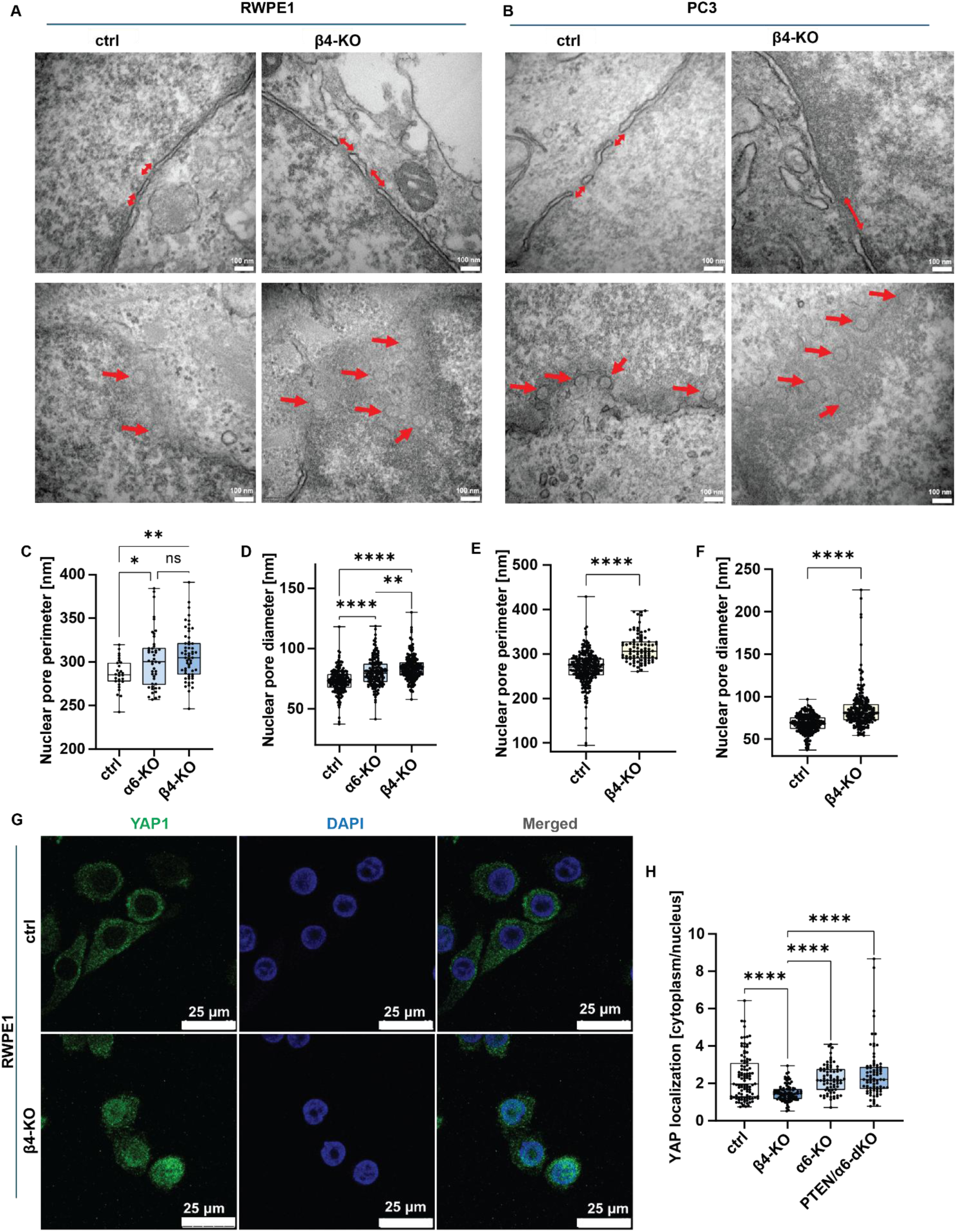
Loss of β4-integrin modulates the nuclear pores. **A-B)** Electron microscopy analysis of RWPE1 (A) and PC3 (B) cells with β4-integrin knock-out. Single red arrows indicate nuclear pores. Analysis was performed using Tecnai Spirit BioTWIN transmission electron microscope. Scale bar, 100 nm. **C-F)** Quantification of nuclear pores perimeter (C and E) and diameter (D and F) in RWPE1 and PC3 cells, respectively. **G)** Localization of the YAP transcription factor in the RWPE1 and β4-KO cells. Cells were stained with anti-YAP1 antibody (green) and DAPI (blue) for nuclei visualization. YAP1 was mainly localized in the cytoplasm in RWPE1 control cells, while it displayed increased nuclear accumulation in RWPE1 β4-KO cells. Images were acquired using Leica SP8X microscopy with a 63x oil objective. Scale bars indicate 25 µm. **H)** Quantification of the YAP localization presented as a ratio of cytoplasmic to nuclear YAP staining. Analysis was performed in LAS X software (1.4.7.28982). Data are presented as box-and-whisker plots (min to max values). Dots represent a number of analyzed nuclear pores or cells.

Together, these observations indicate that β4-depletion is associated with structural changes in nuclear pores and altered subcellular localization of YAP1, supporting a potential role for β4-integrins in maintaining NPC organization and nucleocytoplasmic protein transport.

### H3K27me3-associated heterochromatin is reduced in β4-depleted cells

Transcriptionally inactive heterochromatin is typically enriched at the NL, where it contributes to NE organization and mechanical stability, whereas euchromatin tends to localize more centrally. In cancer, including PCa, the epigenetic landscape often undergoes extensive reorganization, frequently shifting toward a more open chromatin state. To determine whether the NE ruffling observed in β4-deficient cells was associated with altered chromatin organization, we analyzed histone methylation marks characteristic with heterochromatin.

Immunofluorescence analysis revealed that RWPE1 β4-KO cells exhibited significantly reduced levels of H3K9me3 and H3K27me3, markers of constitutive and facultative heterochromatin, respectively, compared with control cells (Supplemental Figure S7A-D). Western blotting analysis further confirmed these findings, demonstrating a pronounced reduction in global H3K27me3 level in β4-KO cells (Supplemental Figure S7E-G). Similar, although more modest, reduction in heterochromatin-associated histone marks were observed in RWPE1 α6-KO cells. In contrast, α6-deficient PC3 cells did not show detectable changes in heterochromatin content compared with parental PC3 cells (Supplemental Figure S7).

These findings indicate that β4-integrin depletion is associated with a loss of facultative heterochromatin and reduced repressive histone methylation. This epigenetic shift is consistent with weakened NL-heterochromatin interactions and further supports a role for β4-integrins in maintaining nuclear organization and chromatin integrity.

## DISCUSSION

Disassembly of HDs and the loss of β4-integrin expression are well-documented events during PCa progression and have been shown to actively promote tumorigenesis. Here, our study uncovers a previously unappreciated function of β4-integrins in preserving nuclear architecture and mechanical integrity. We demonstrate that loss of β4-integrin expression disrupts CK5-based intermediate filament (IF) cytoskeleton, weakens IF-nuclear envelope (NE) coupling, and induces NE ruffling and nuclear softening. These mechanical alterations were accompanied by a lamin isoform shift, characterized by reduced lamin A/C levels and increased lamin B1 expression, a characteristic of aggressive and mechanically adaptable cancers. Proximity-based proteomics revealed that β4-integrins interact with NPC proteins. Intriguingly, β4-integrin depletion affected nuclear pore structure and function. Collectively, our findings position β4-integrins as critical regulators of nuclear homeostasis whose loss promotes nuclear adaptability, genome integrity and malignant progression.

β4-integrin is a transmembrane protein synthesized in the ER, where it forms a heterodimer with the α6-subunit. α6β4-heterodimer then undergoes biosynthetic trafficking from the ER through the Golgi and trans-Golgi network to the plasma membrane to assemble HDs. Physiologically, HD turnover is governed by phosphorylation of the β4-integrin cytoplasmic tail by kinases such as PKC, Src and Erk, which disrupts β4-plectin binding and release keratin filaments from the basal membrane(13,60). Importantly, β4-integrins are expressed in stoichiometric excess relative to α6-integrins, suggesting that β4-subunits could have also α6-independent functions(24). In the absence of α6, unpaired β4-subunits are thought to be largely retained in the ER(61). However, despite this predominantly intracellular retention, β4-integrins can interact with multiple protein partners in α6-KO cells(25). Our proximity-based proteomic analyses in the current study support this view by revealing an α6-independent β4-integrin interactome enriched for NPC components, particularly upon HD disassembly. These findings raise the possibility that β4-integrins perform non-canonical intracellular functions, potentially within the ER-NE continuum, that contribute to nuclear organization and function.

Functionally, loss of β4-integrin leads to nuclear softening and altered NL composition, changes that may promote tumorigenesis through several, not mutually exclusive, mechanisms. First, nuclear softening may render nuclei more deformable, facilitating invasive migration through confined spaces within dense tumor tissues(62,63). Second, a softer nucleus may partially buffer nuclei and preserve genomic integrity against acute compressive stresses encountered *in vivo*(64,65). Third, the observed alterations in NPCs could deregulate nucleocytoplasmic transport, as exemplified by the YAP1 transcription factor, thereby enhancing nuclear entry of oncogenic transcriptional regulators and supporting tumorigenic transcriptional programs(66). This finding is particularly intriguing, as YAP1 nuclear localization is typically associated with increased cytoskeletal tension and nuclear stiffness. The apparent uncoupling of YAP1 nuclear entry from cytoskeletal tension in β4-deficient cells suggests that aberrant NPC function, altered NL composition, or compromised NE integrity may override canonical mechanotransduction pathways. Consistent with this notion, Roca-Cusachs and coworkers reported that HD-anchored keratin cytoskeleton decouples YAP nuclear transport from substrate rigidity, thereby protecting the nucleus from mechanical deformation(45). In that study, function-blocking β4-integrin antibodies or expression of a plectin-binding-deficient β4-integrin mutant restored nuclear YAP transport on rigid substrates and induced changes in nuclear morphology. Importantly, elegant prior work has demonstrated that force transmission through the LINC complex can deform the NE, widen nuclear pores, and enhance YAP translocation independently of global cytoskeletal stiffening(67). Whether β4-integrins influence NPC function through direct interactions with nucleoporins, indirect modulation of the LINC complex, or altered IF-mediated force transmission at NE remains an important question for future investigation.

Beyond nucleocytoplasmic transport, β4-integrin interactions with nucleoporins such as TPR may also influence higher-order chromatin organization. TPR is a key regulator of perinuclear chromatin architecture, contributing to chromatin exclusion zones and the maintenance of transcriptionally permissive domains near nuclear pores(68). In line with this, we observed reduced levels of heterochromatin-associated histone marks, particularly H3K27me3, in β4-deficient cells. Disruption of NE structure and NPC organization may therefore release heterochromatin from the nuclear periphery, increase chromatin accessibility and sensitize cells to oncogenic transcriptional reprogramming(64,69). Such chromatin remodeling could synergize with aberrant nucleocytoplasmic transport to reinforce malignant gene expression programs. Notably, H3K27me3 levels were increased in cells expressing the plectin-binding–deficient β4-integrin mutant (R1281W)(45), whereas β4-integrin loss reduced heterochromatin levels. This divergence indicates that β4-integrins regulate chromatin state through mechanisms that are not strictly dependent on plectin-mediated cytoskeletal anchorage and may involve distinct nuclear regulatory functions.

Importantly, our mechanistic findings are strongly supported by clinical data. Reduced β4-integrin expression in patient samples correlates with nuclear deformation, altered lamin expression, and poor clinical outcome, including metastasis and PCa-specific mortality. Notably, these associations are evident not only in tumor tissue but also in phenotypically normal epithelium, suggesting that β4-integrin loss may represent an early event that predisposes cells to malignant transformation by weakening nuclear integrity. Assessment of β4-integrin status, together with nuclear features and lamin expression patterns, may improve risk stratification beyond current histopathological criteria by identifying biomechanically unstable tumors predisposed to aggressive progression.

Taken together, our data identifies β4-integrins as key suppressors of malignant progression through their previously unrecognized role in maintaining nuclear architecture, mechanical stability, and genome integrity. By linking integrin-mediated adhesion, IF organization, nuclear mechanics, NPC function, and chromatin regulation, we establish a new conceptual framework that connects cell-matrix adhesion biology to nuclear homeostasis in prostate cancer. These findings provide a mechanistic rationale for the selective loss of β4-integrins during PCa progression and suggest that restoration of nuclear integrity pathways, rather than adhesion alone, may represent a promising therapeutic avenue.

## ACKNOWLEDGEMENTS

We thank Riitta Jokela for overall expert technical assistance, Mari Sujala for expert technical assistance at Biocenter Oulu Virus Core Laboratory, Dr. Veli-Pekka Ronkainen and Dr Virpi Glumoff for expert assistance in FACS. Biocenter Finland, University of Oulu are acknowledged for contributing research infrastructure services. We are very grateful to Professor Barbara Lipinska from the Department of General and Medical Biochemistry (University of Gdansk, Faculty of Biology) for her many helpful suggestions and comments.

This work was funded by Research Council Finland profiling program (grant #311934) to AM, Jane and Aatos Erkko foundation (190046/AM, GHW), Sigrid Jusélius Foundation (GHW), Cancer Foundation Finland (GHW, 61-6158/AM), Biocenter Oulu spearhead funding to GW and AM, Magnus Ehrnrooth foundation to AM, University of Oulu and RCF Profi6 Fibrobesity programme #336449 to AM, National Science Centre, Poland - grant UMO-2023/51/D/NZ2/00561 to TW, grant UMO-2020/39/D/NZ3/00882 to PN, University of Gdansk - Support program for Science Leaders 2026 (No. 539-D010-B290-26) to TW, the Finnish Cultural Foundation (00241065/AS), Cancer Foundation Finland sr (5029/AS), University of Oulu Scholarship Foundation (20250150/AS) and Orion Research Foundation sr (15-13835-27/AS) to AS. AN and AA acknowledge partial funding from the National Science Foundation (NSF, Grant No. 2422340 and 2119949) and the National Institute of Health (1R01 HL162822-01A1). AN and AA acknowledge the Institute of Critical Technologies and Science (ICTAS) and Macromolecules Innovative Institute (MII) at Virginia Tech for supporting this study.

## DATA AVAILABILITY

The mass spectrometry proteomics data generated in this study have been deposited to the MassIVE repository under accession number MSV000100146. The RNA-sequencing data have been deposited in the European Nucleotide Archive (ENA) under accession number PRJEB105719.

## AUTHOR CONTRIBUTION

Conceptualization: TW(1,2), AM(1); Methodology: AA(4), CR(7,8,9), CFN(9), AN(4); Validation: AS(1), RW(2), PN(3), MME(1), KB(6), SK(1), XY(1), TW(1,2); Formal Analysis: RW(2), QZ(1), XL(11), MV(13), TW(1,2); Investigation: AS(1), RW(2), PN(3), MME(1), TW(1,2); Resources: AA(4), MR(5), KB(6), AA(10), MV(10), KDR(12), EB(12), AN(4); Writing — Original Draft: TW(1,2), AM(1); Writing — Review and Editing: AS(1), RW(2), PN(3), MR(5), CF(7,8,9), CFN(9), XL(11), AN(4), GHW(1,14,15), TW(1,2), AM(1); Visualization: AS(1), RW(2), MR(5), TW(1,2); Supervision: TW(1,2), AM(1); Funding Acquisition: TW(1,2), AM(1).

## ONLINE SUPPLEMENTAL MATERIAL

Online supplemental material includes seven (7) supplemental figures: **Figure S1**. RNA-Seq analysis of RWPE1 PTEN-KO and PTEN/α6-dKO cells. **Figure S2.** Loss of β4-integrins affects nuclear morphology in PC3 cells. **Figure S3.** Electron microscopy analysis of DU145 β4-KO cells. **Figure S4.** Changes of β4-integrin levels affect nuclear morphology of breast and prostate cells. **Figure S5**. β4-integrin interacts with proteins regulating cell adhesion and nuclei function. **Figure S6**. HD disassembly in α6-KO cells drives a shift in the β4-integrin interactome toward nuclear-associated proteins. **Figure S7.** Heterochromatin-associated H3K27me3 is reduced in β4-deficient cells. **Figure S8.** Prognostic value of the nuclear morphology and clinical parameters of patients from TMA.

Three (3) supplemental tables: **Table S1.** Complete processed datasets underlying the transcriptomic (Figure 1) and proteomic (Figure 7) analyses. **Table S2.** List of antibodies used in this study. **Table S3.** List of qPCR primers used in this study.

Four (4) supplemental videos: **Video S1**. Representative time-lapse images of PC3 control cells migrating through the constrictive channels of 3D-fabricated cell confinement device. **Video S2**. Representative time-lapse images of PC3 control cells migrating through the non-constrictive channels of 3D-fabricated cell confinement device. **Video S3**. Representative time-lapse images of PC3 β4-KO cells migrating through the constrictive channels of 3D-fabricated cell confinement device. **Video S4**. Representative time-lapse images of PC3 β4-KO cells migrating through the non-constrictive channels of 3D-fabricated cell confinement device.

