## Supplemental data for "β4-integrins safeguard nuclear mechanics to suppress prostate cancer progression"

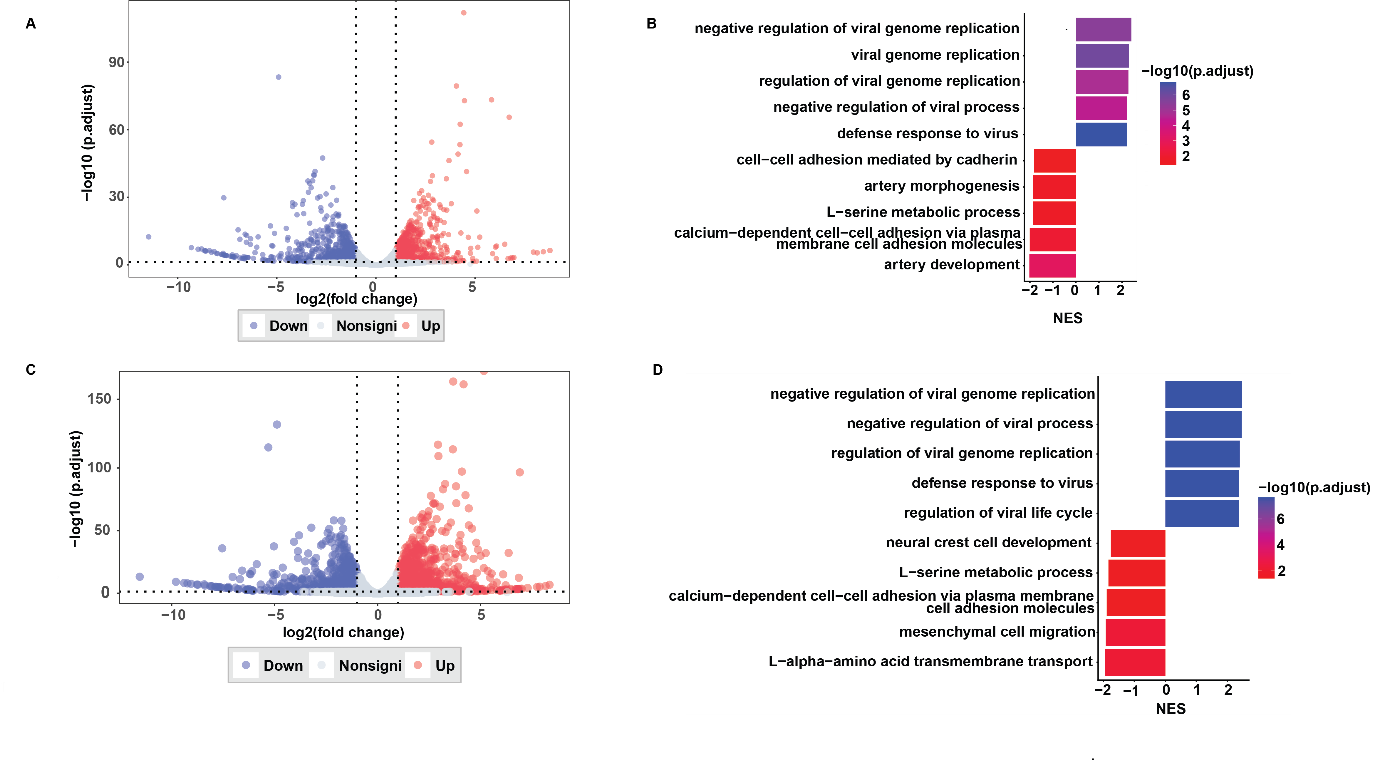
***Supplemental Figure S1****.* ***RNA-Seq analysis of RWPE1 PTEN-KO and PTEN/α6-dKO cells.***

***A)*** *Volcano plot of differentially expressed genes (DEGs) in RWPE1 PTEN/α6-dKO versus control cells.*

***B)*** *GSEA of genes ranked by log_2_ fold change in RWPE1 PTEN/α6-dKO cells.*

***C)*** *Volcano plot of DEGs in RWPE1 PTEN-KO vs. control. The x-axis represents log₂ (fold change) and the y-axis represents -log₁₀ (adjusted p-value). Red and blue dots indicate significantly up- and down-regulated genes, respectively, while grey dots represent non-significant genes. Significance was defined as |log₂ fold change| > 1 and adjusted p < 0.05.*

***D)*** *GSEA of genes ranked by log_2_ fold change in RWPE1 PTEN-KO cells. The x-axis shows the normalized enrichment score (NES), and the y-axis lists the top 5 significantly up- and down-regulated biological processes. The color gradient corresponds to -log₁₀ (adjusted p-value).*

*Full list of enriched pathways shown in panels B and D is provided in Supplemental Table S1.*

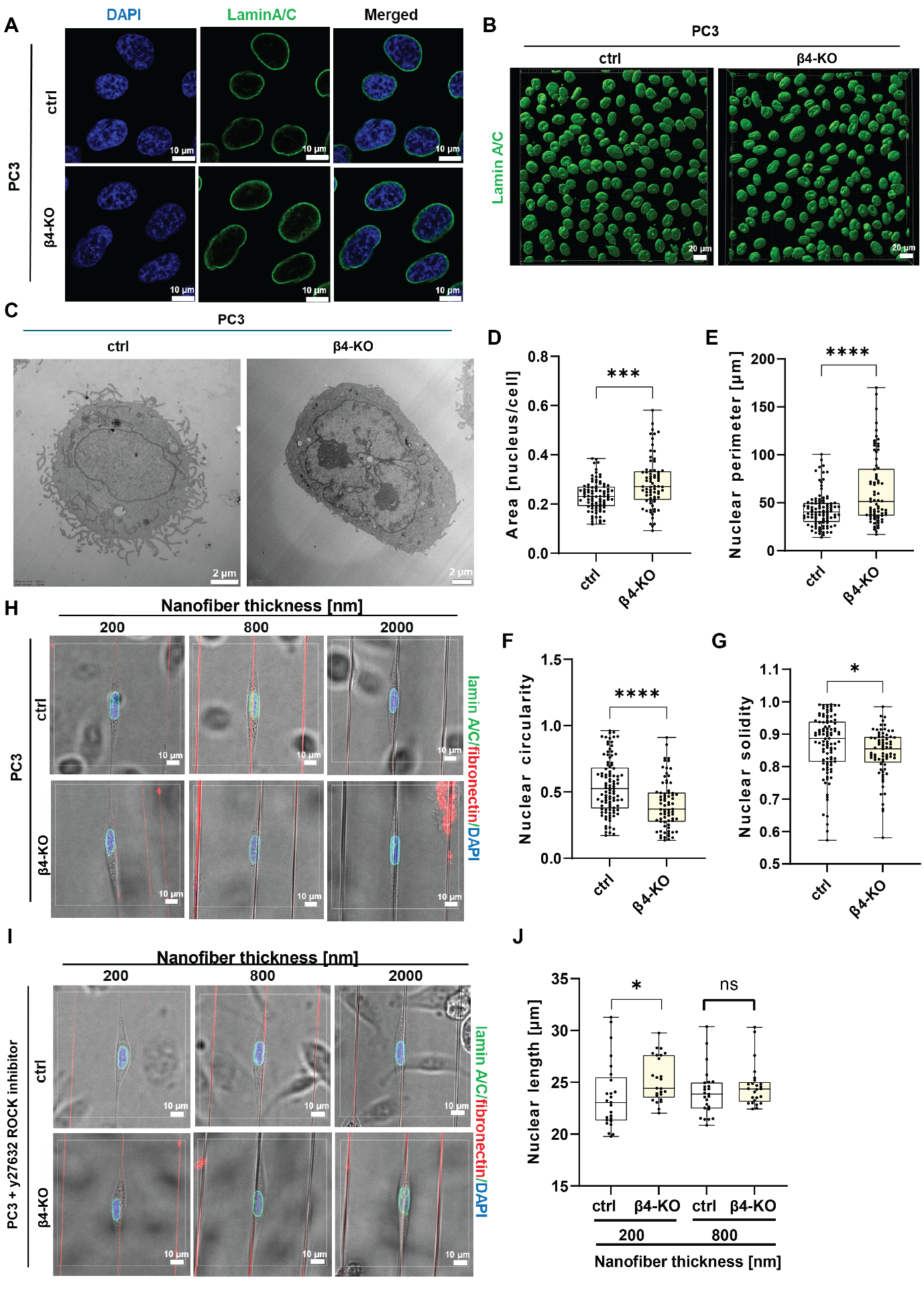
**Supplemental Figure S2**. **Loss of β4-integrins affects nuclear morphology in PC3 cells**.

***A)*** *Nuclear morphology of PC3 and PC3 β4-KO cell lines visualized by lamin A/C (green) and DAPI (blue) staining. Images were acquired using a Leica SP8 Falcon confocal microscope with a 40x water objective. Scale bars represent 10 µm.*

***B)*** *3D-rendered nuclear morphology of PC3 control and β4-KO cells based on lamin A/C staining. Analysis was performed using IMARIS software. Scale bars represent 20 µm.*

***C)*** *Electron microscopy analysis of nuclei of PC3 control and β4-KO cells. Analysis was performed using Tecnai Spirit BioTWIN transmission electron microscope. Scale bars represent 20 µm.*

***D-G)*** *Quantification of nuclear morphology from electron microscopy images. PC3 β4-KO cells exhibited increased nuclear area (D) and perimeter (E), and reduced circularity (F) and solidity (G). Analysis was performed using ImageJ software.*

***H-I)*** *Merged brightfield images with DAPI (blue), lamin (green) and fibronectin-coated nanofibers (red) showing nuclei of PC3 control and β4-KO cells. Cells were cultured either in complete F12K medium (H) or treated with Y-27632 ROCK inhibitor (I). Brightfield imaging was used to visualize cell shape. Images were acquired using Leica SP8 Falcon confocal microscope with a 40x water objective. Scale bars represent 10 µm.*

***J)*** *Quantification of nuclear length in cells grown on nanofiber of different diameters. Dots represent individual analyzed cells or nuclei. Data are presented as box-and-whisker plots (min to max values).*

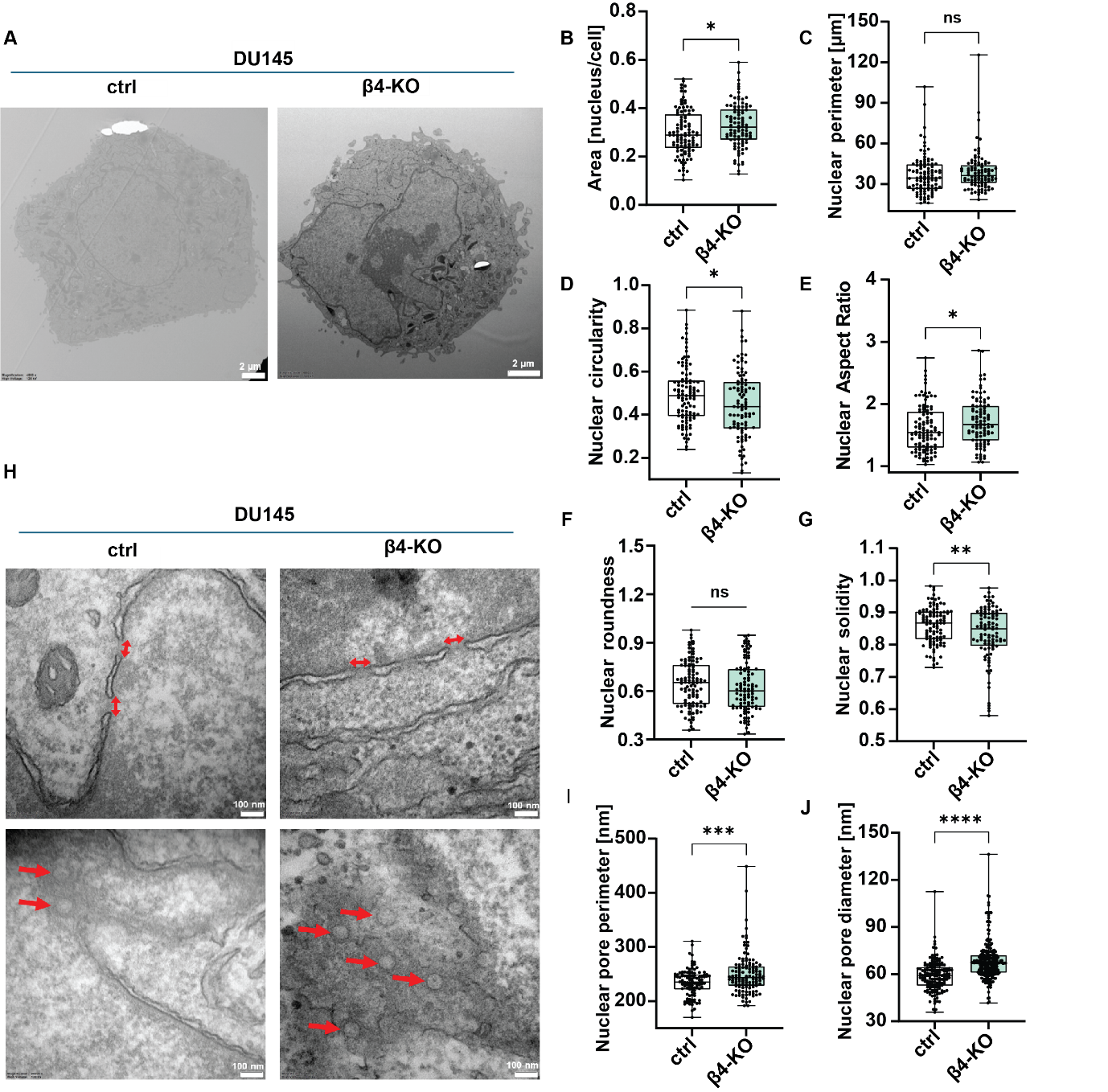
**Supplemental Figure S3**. ***Electron microscopy analysis of DU145 β4-KO cells****.*

***A)*** *Electron microscopy images of nuclei in DU145 control and β4-KO cells. Analysis was performed using Tecnai Spirit BioTWIN transmission electron microscope.*

***B-G)*** *Quantification of nuclear morphology based on electron microscopy images. Nuclear area (B), perimeter (C), circularity (D), aspect ratio (E), roundness (F), and solidity (G) were measured using ImageJ software. Dots represent individual analyzed nuclei.* *Data are presented as box-and-whisker plots (min to max values).*

***H)*** *Electron microscopy analysis of nuclear pores in DU145 control and β4-KO cells. Single red arrows indicate nuclear pores. Analysis was performed using Tecnai Spirit BioTWIN transmission electron microscope. Scale bar, 100 nm.*

***I-J)*** *Quantification of nuclear pore perimeter (I) and diameter (J) in DU145 cells. Dots represent individual analyzed nuclear pores.*

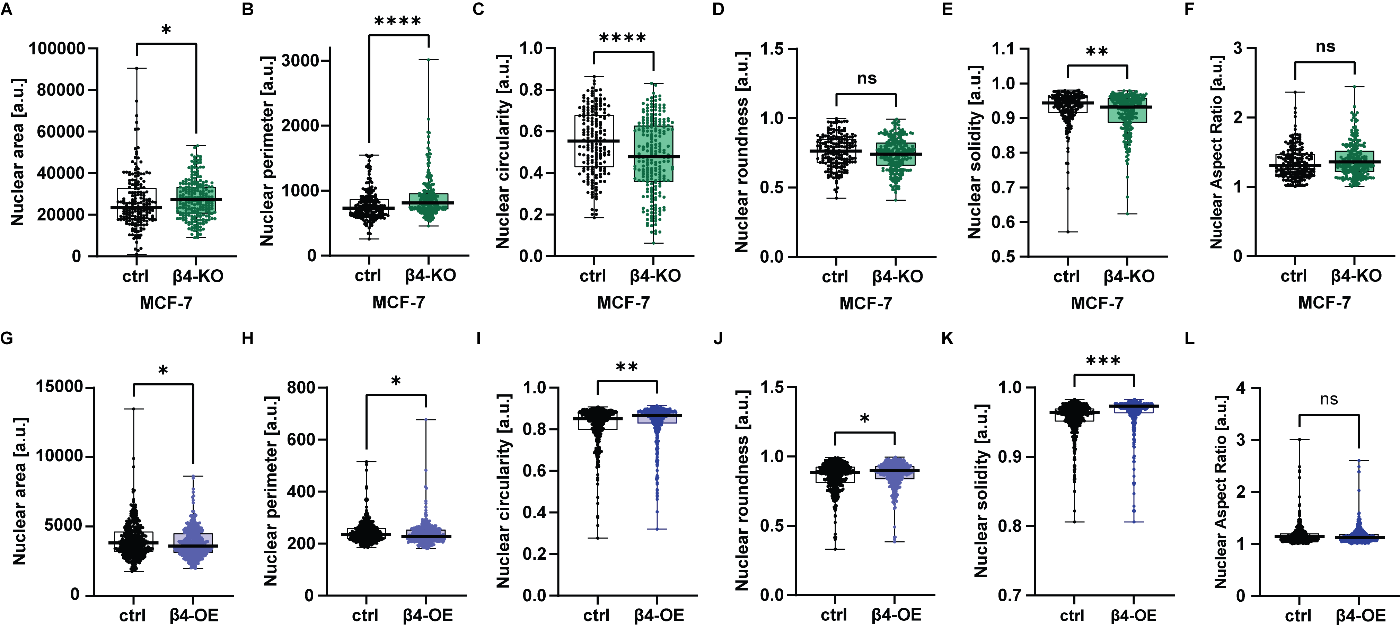

**Supplemental Figure S4**. **Changes of β4-integrin levels affect nuclear morphology of breast and prostate cells.** *Quantification of nuclei from MCF7 breast cancer cells with β4-integrin depletion* ***(A-F)*** *and RWPE1 with overexpression of β4-integrin* ***(G-L).*** *Nuclear area* ***(A, G)****, perimeter* ***(B, H)****, circularity* ***(C, I)****, roundness* ***(D, J)****, solidity* ***(E, K)*** *and aspect ratio* ***(F, L)*** *were measured on confocal microscopy images using ImageJ software. Nuclei were stained by using DAPI. Dots represent individual analyzed nuclei.* *Data are presented as box-and-whisker plots (min to max values).*

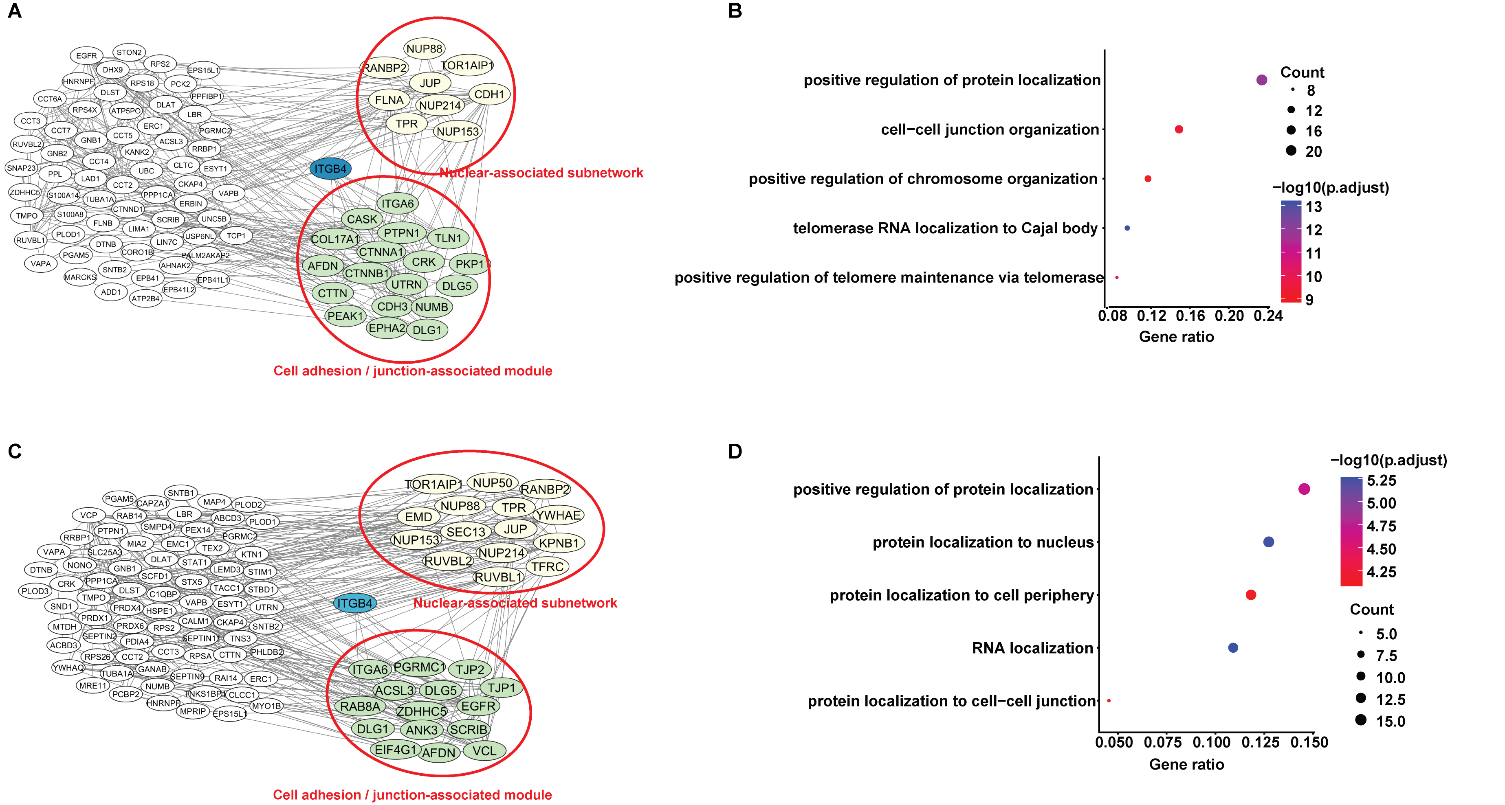

**Supplemental Figure S5**. ***β4-integrin interacts with proteins* regulating cell adhesion and nuclei function.**

**A, C)** *Protein-protein interaction (PPI) maps of β4-integrin proximity partners in RWPE1 (A) and PC3 (C) cells overexpressing β4-integrin-BirA. Candidate interactors were identified using the MiST algorithm as described in the Methods section and visualized using Cytoscape (version 3.10.3). Full list of candidate interactors is provided in Supplemental Table S1.*

***B, D****) GO enrichment analysis of HICs identified using the MIST algorithm. The x-axis shows the Rich factor and the y-axis lists the top enriched biological processes. Dot size corresponds to the number of proteins in each term and color indicates -log₁₀(adjusted p-value).*

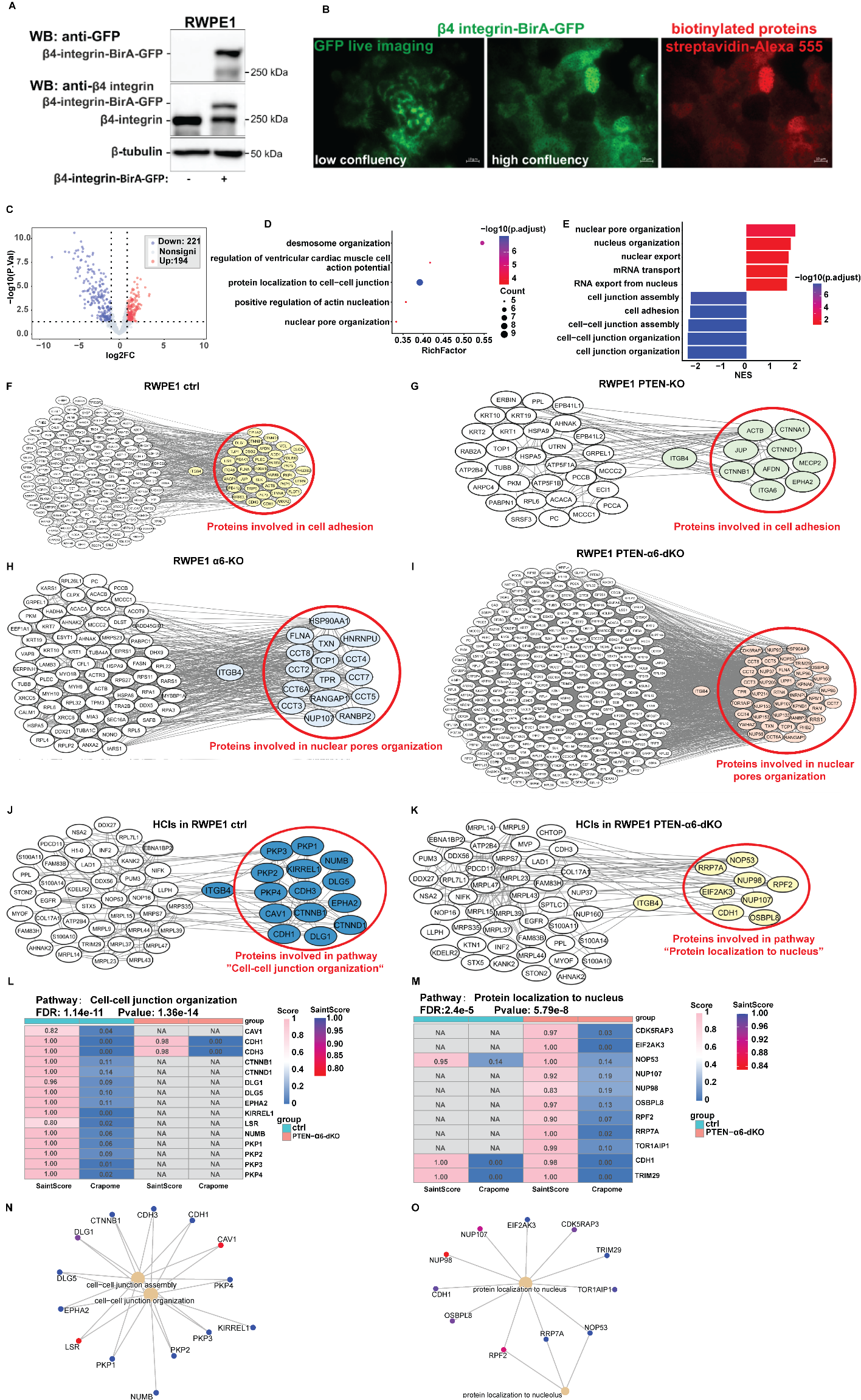

**Supplemental Figure S6. *HD disassembly in α6-KO cells drives a shift in the β4-integrin interactome toward nuclear-associated proteins.***

***A-B)*** *Generation of RWPE1 cells with “knock-in” of BirA-GFP into ITGB4 sequence. Modified cells were analyzed by western blotting (A) and by TIRF microscopy (B) to verify the proper cellular localization.*

***C)*** *Volcano plot of differentially expressed proteins (DEPs) identified by BioID of endogenous β4-integrin in RWPE1 α6-KO compared with control cells. A total of 194 proteins were enriched and 221 were depleted. The x-axis shows log₂ fold change and the y-axis shows -log₁₀(adjusted p-value). Significance was defined as |log₂ fold-change| > 1 and p-value less than 0.05.*

***D-E)*** *GO enrichment (B) and GSEA enrichment (C) analyses of proteins ranked by log₂ fold-change in α6-KO cells. For GO analysis, the x-axis represents the Rich factor and the y-axis lists the top five enriched biological processes. Dot size indicates the number of proteins per term and color indicates -log₁₀ (adjusted p-value). For GSEA, the x-axis shows the normalized enrichment score (NES) and the y-axis shows the top enriched pathways.*

***F-I)*** *Protein-protein interaction (PPI) maps of β4-integrin proximity partners in RWPE1 control (F), PTEN-KO (G), α6-KO (H) and PTEN-α6-dKO (I) cells, visualized using Cytoscape (version 3.10.3). Candidate interactors were identified using the MiST algorithm as described in the Methods section. Full list of candidate interactors is provided in Supplemental Table S1.*

***J-K)*** *PPI maps showing high-confidence interaction (HCI) of β4-integrin in RWPE1 control (J) and PTEN-α6-dKO (K) cells identified using SAINT analysis. Full list of HCI is provided in Supplemental Table S1.*

***L-M)*** *Heatmaps showing β4-integrin HCIs in the top-ranked pathways derived from over-representation analysis (WebGestalt http://www.webgestalt.org) of β4-integrin HCIs in RWPE1 control (L) and PTEN-α6-dKO (M) cells.*

***N-O)*** *Enrichment network plots of β4-integrin HCIs in RWPE1 control (N) and PTEN-α6-dKO (O) cells highlighting top enriched pathways, generated using the cnetplot function in the clusterProfiler package (version: 4.14.6).*

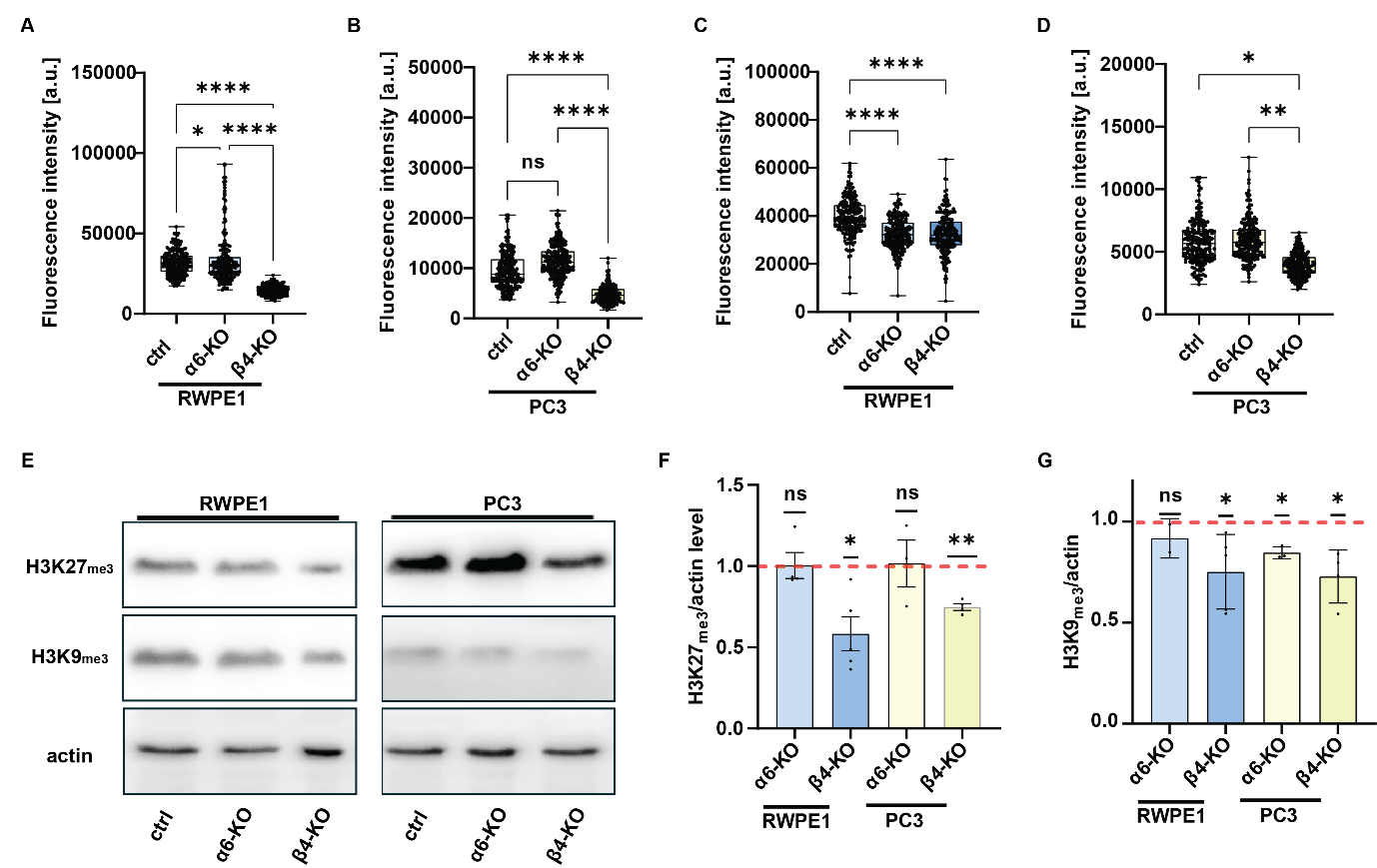
**Supplemental Figure S7**. **Heterochromatin-associated H3K27me3 is reduced in β4-deficient cells.**

#### **A-D)** β4- and α6-depleted variants of RWPE1 (A, C) and PC3 (B, D) cells were stained for the heterochromatin marks H3K27me3 (A, B) and H3K9me3 (C, D). Fluorescence intensity was quantified using LAS X software. Dots represent individual analyzed cells. Reduced levels of both heterochromatin marks were consistently observed in β4-deficient cells. The effects in α6-KO cells were less pronounced, suggesting that loss of β4-integrins is required to induce robust nuclear and heterochromatin phenotypes.

#### **E)** Western blot analysis showing decreased levels of H3K27me3 and H3K9me3 in β4-integrin depleted RWPE1 and PC3 cells.

#### **F-G)** Quantification of western blot results for H3K27me3 (F) and H3K9me3 (G). Red dashed lines indicate normalized levels of these histone marks in parental control cells. Dots represent biological replicates. Data are presented as mean ± min/max values.

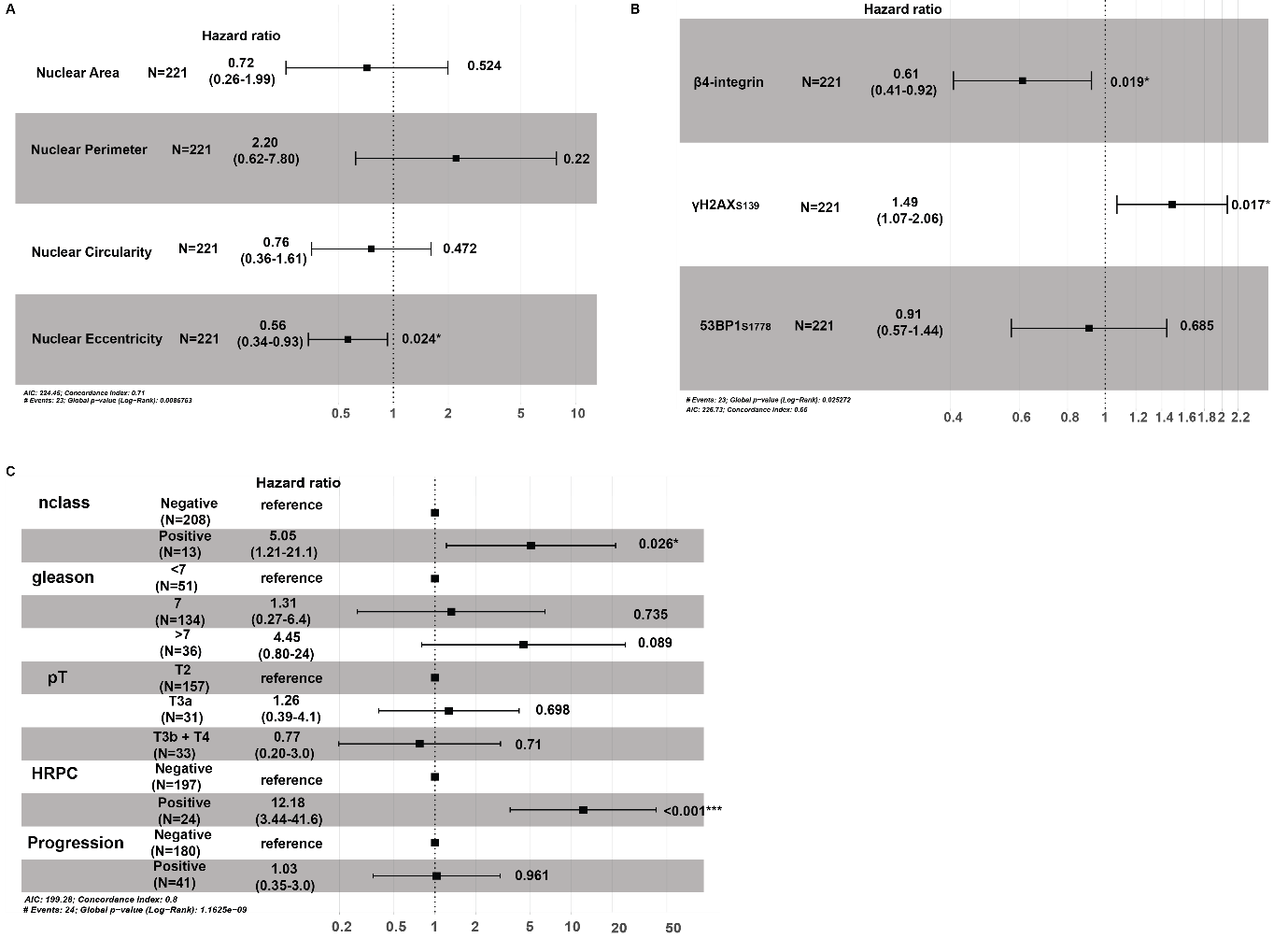
**Supplemental Figure S8. P*rognostic value of the nuclear morphology and clinical parameters of patients from TMA.***

***A-C)*** Forest plots displaying proportional hazards analyses estimates of nuclear morphology parameters including nucleus area, perimeter, circularity, and eccentricity (A), β4-integrin, pH2AX_S139_ and p53BP1_S1778_ levels (B) and clinical covariates: lymph node involvement, Gleason score, pathological stage, hormone-refractory status and PCa progression (C) of 221 patients with PCa. Hazard ratio (HR), representing the relative change in risk associated with the variable of interest, are shown with 95% confidence intervals; HRs > 1 indicate increased risk, whereas HRs < 1 indicate reduced risk. Statistical significance was assessed using Cox proportional hazards regression. *P < 0.05; ***P < 0.001.

#### Supplemental Table S2. List of antibodies used in this study.

| Name | Type | Company | Cat. No | Dilution for WB/PLA | Dilution for ICC/IHC |
| --- | --- | --- | --- | --- | --- |
| anti-α6-integrin | polyclonal | Sigma Aldrich/Merck | HPA012696 | 1:2000/1:200 |  |
| anti-β-actin | monoclonal | Sigma-Aldrich/Merck | A3854 | 1:10000 |  |
| anti-β4-integrin | monoclonal | Abcam | ab29042 | 1:500/1:50 | 1:50/1:50 |
| anti-β4-integrin | monoclonal | AbClonal | A24995 | 1:1000 |  |
| anti-cytokeratin 5 | polyclonal | Proteintech | 28506-1-AP |  | 1:100 |
| anti-fibronectin | polyclonal | Sigma-Aldrich/Merck | F3648 |  | 1:500 |
| anti-GAPDH | monoclonal | Cell Signaling Technology | 5174 | 1:10000 |  |
| anti-LaminA/C | monoclonal | Santa Cruz Biotechnology | sc-376248 |  | 1:1000 |
| anti-Lamin A/C | monoclonal | Abcam | ab215495 | 1:5000 | 1:2000 |
| anti-Lamin A/C | monoclonal | Abcam | ab320906 | 1:5000 | 1:2000 |
| anti-Lamin B1 | polyclonal | Abcam | ab16048 | 1:2000 |  |
| anti-Lamin B1 | monoclonal | Abcam | ab194108 |  | 1:2000 |
| anti-NUP98 | monoclonal | AbClonal | A22743 | 1:1000/1:100 |  |
| anti-NUP98 | monoclonal | Santa Cruz Biotechnology | sc74578 | 1:1000/1:100 |  |
| anti-TPR | monoclonal | Santa Cruz Biotechnology | sc271565 | -/1:100 |  |
| anti-TPR | polyclonal | Merck | HPA024336 | 1:2000 | 1:50 |
| anti-tri-methyl-histone H3 (Lys9) | monoclonal | Cell Signaling Technology | 13969 | 1:1000 | 1:1000 |
| anti-tri-methyl-histone H3 (Lys27) | monoclonal | Cell Signaling Technology | 9733 | 1:1000 | 1:1000 |
| anti-YAP | monoclonal | AbClonal | A21216 |  | 1:500 |
| anti-53BP1 | polyclonal | Novus Biologicals | NB-100-305 |  | 1:500 |
| anti-53BP1 | polyclonal | AbClonal | A5757 | 1:5000 |  |

#### Supplemental Table S3. List of qPCR primers used in this study.

| Gene | Sequence 5’-3’ | Predicted product size [bp] |
| --- | --- | --- |
| *MAD2L1* | F: GCCGAAATCGTGGCCGAG  R: GTTACAAGCAAGGTGAGTCCGT | 119 |
| *INCENP* | F: GAGCTGATGCCCAAAACACCT  R: TGCGGGATAACCTTCTCCTGAT | 106 |
| WRAP53 | F: CCAGCTCTTCTGTGGCTTCAAC  R: AGGCTATGCAGGAGATGATGCC | 128 |
| GAPDH | F: GTCTCCTCTGACTTCAACAGCG  R: ACCACCCTGTTGCTGTAGCCAA | 131 |
